# PKA Activity-Driven Modulation of Bidirectional Long-Distance transport of Lysosomal vesicles During Synapse Maintenance

**DOI:** 10.1101/2024.06.28.601272

**Authors:** Kerriann K. Badal, Yibo Zhao, Bindu L Raveendra, Sebastian Lozano-Villada, Kyle E. Miller, Sathyanarayanan V. Puthanveettil

**Affiliations:** Department of Neuroscience, The Herbert Wertheim UF Scripps Institute for Biomedical Innovation & Technology, 130 Scripps Way, Jupiter, FL 33458, USA; Integrative Biology PhD Program, Charles E. Schmidt College of Science, Florida Atlantic University, Jupiter, FL 33458, USA; Harriet L. Wilkes Honors College, Florida Atlantic University, 5353 Parkside Drive, Jupiter, FL 33458, USA; Department of Integrative Biology, Michigan State University, East Lansing, MI 48824, USA

**Keywords:** Synapse formation, synapse maintenance, plasticity, long-distance transport, gene expression, lysosome related organelles, mitochondria, Protein kinase A

## Abstract

The bidirectional long-distance transport of organelles is crucial for cell body-synapse communication. However, the mechanisms by which this transport is modulated for synapse formation, maintenance, and plasticity are not fully understood. Here, we demonstrate through quantitative analyses that maintaining sensory neuron-motor neuron synapses in the *Aplysia* gill-siphon withdrawal reflex is linked to a sustained reduction in the retrograde transport of lysosomal vesicles in sensory neurons. Interestingly, while mitochondrial transport in the anterograde direction increases within 12 hours of synapse formation, the reduction in lysosomal vesicle retrograde transport appears three days after synapse formation. Moreover, we find that formation of new synapses during learning induced by neuromodulatory neurotransmitter serotonin further reduces lysosomal vesicle transport within 24 hours, whereas mitochondrial transport increases in the anterograde direction within one hour of exposure. Pharmacological inhibition of several signaling pathways pinpoints PKA as a key regulator of retrograde transport of lysosomal vesicles during synapse maintenance. These results demonstrate that synapse formation leads to organelle-specific and direction specific enduring changes in long-distance transport, offering insights into the mechanisms underlying synapse maintenance and plasticity.

## Introduction

Synapse formation involves multiple specializations requiring the coordinated regulation of molecular and biochemical pathways in pre- and postsynaptic neurons (Shen and Scheiffele, 2010; West and Greenberg, 2011; Sudhof, 2018). In presynaptic neurons, these include active zone formation, vesicle clustering, neurotransmitter synthesis and release, and mitochondrial enrichment (Lee et al., 2008; Tang and Zucker, 1997; Brodin et al., 1999; Blaustein et al., 1978). In postsynaptic neurons, they include post-synaptic densities and neurotransmitter receptor clustering (Bourne and Harris, 2008; Sheng, 2001). Additionally, specialized cell adhesion complexes, transsynaptic signaling proteins, and growth factors are required (Shen and Scheiffele, 2010; West and Greenberg, 2011; Dickins and Salinas, 2013; Sudhof, 2018; Missler et al., 2012). Retrograde signaling from postsynaptic neurons also modulates presynaptic signaling (Williams, 1996; Sastry et al., 1988; Tao and Poo, 2001). While these studies illuminate synapse formation, little is known about mechanisms by which synapses are maintained (Lin and Koleske, 2010; Heckman and Doe, 2021).

Recent research using models like Drosophila, C. elegans, and mice has identified few key players in synapse maintenance, such as ATAT-2 tubulin acetyl transferase, the RPM-1 signaling hub (Borgen et al., 2019), IP3 signaling (James et al., 2019), Shank 3 phosphorylation (Wu et al., 2022), polarity proteins (Voglewede and Zhang, 2022), calcium channels (Cunningham et al., 2022), Piccolo (Garner et al., 2023), and cell adhesion molecules (Nabavi and Hiesinger, 2023). Together these findings describe local mechanisms governing synapse maintenance.

Given the highly polarized architecture of neurons, with the soma communicating with synapses through axons and extensive dendritic arbors, it is likely that soma-synapse communication is essential for synapse maintenance. Several studies, including our own, have demonstrated that long-distance delivery of gene products from the soma to the synapse via molecular motor proteins along microtubule tracks (Yamada et al., 1970) ensures proper synapse function and plasticity (Puthanveettil et al., 2008; Puthanveettil et al., 2013; Liu et al., 2014; Swarnkar et al., 2018; Swarnkar et al., 2021; Joseph et al., 2021; Panayotis et al., 2015; Guedes-Dias & Holzbaur, 2019). Kinesin mediates anterograde transport to deliver proteins, RNAs, and organelles to synapses (Saxton & Hollenbeck, 2012; Guillaud et al., 2020; Maday et al., 2014), while dynein/dynactin-facilitated retrograde transport is crucial for cellular signaling, material recycling, and neuronal homeostasis (Schwartz, 1979; Yamashita, 2019; Guedes-Dias & Holzbaur, 2019). However, whether and how this bidirectional transport is modulated to maintain synapse function remains unclear, representing a critical gap in our understanding of neural circuit function and plasticity.

To assess the role of long-distance transport in synapse maintenance, we utilized the Aplysia gill-siphon withdrawal reflex model. Unlike vertebrate neurons that form autapses, Aplysia sensory neurons (SNs) and motor neurons (MNs) form synapses exclusively with specific neuron types (Kandel, 2001; Bailey et al., 2015; Glanzman, 2006; Hawkins et al., 2006). In the presence of L7MN, a target postsynaptic MN in the abdominal ganglia, SNs form functional synapses that remodel and grow in response to neuronal stimulation (Miniaci et al., 2008; Upreti et al., 2019). We recently reported that maintaining SN-L7MN synapses involves a bidirectional increase in mitochondrial transport flux, with anterograde transport significantly enhanced within 24 hours of synapse formation (Badal et al., 2029). Nonetheless, it is not yet clear whether this increase is global or cargo-specific, and the regulatory mechanisms of transport during synapse maintenance and plasticity remain to be understood.

Using the SN-L7MN cell culture system, we examined whether synapse maintenance affects transport in a cargo-specific manner and whether synapse formation and plasticity uniquely modulate different cargos. We imaged anterograde and retrograde transport of lysosomal vesicles during synapse maintenance, formation, and plasticity. Lysosomal vesicles (LV), a group of acidic organelles involved in the endolysosomal system, autophagy, and macromolecule recycling, are essential for neuronal connectivity, sensory-motor functions, mood, learning, memory, and aging (Delevoye et al., 2019; Ferguson, 2018; Kornfeld, 1989; Roney et al., 2022; Dell’Angelica et al., 2000; Luzio et al., 2014; Bezprozvanny et al., 2013; Azpurua et al., 2015; Ding et al., 2022). Defects in lysosomal vesicles axonal transport and signaling impair axonal homeostasis and are linked to neurodegenerative diseases (Roney et al., 2021; Lee et al., 2011; Roney et al., 2022). Our quantitative live imaging of lysosomal vesicle transport revealed direction-specific modulation, with a significant reduction in retrograde transport observed 72 hours after synapse formation and further reduction within 24 hours after induction of plasticity. Pharmacological inhibition studies suggest that persistent changes in transcription and PKA activity underlie the modulation of lysosomal vesicle transport.

## Results

### Reduction in the long-distance retrograde transport of lysosomal vesicles during the maintenance of SN-L7MN synapses

To examine whether synapse maintenance is associated with a general enhancement in the bidirectional transport of organelles, such as mitochondria in presynaptic sensory neurons (SNs) (Badal et al., 2019), or with organelle-specific and direction-specific modulation of transport, we imaged long-distance transport of lysosomal vesicles. In SN and postsynaptic L7 motor neuron (L7MN) co-cultures (SN-L7MN), synapses begin to form within 6 hours of plating. In these cultures, the excitatory postsynaptic potentials (EPSPs), a measure of synaptic communication, increase until three days after in vitro culture (DIV3) and then remain stable. Therefore, SN-L7MN cultures that are more than three days old are considered to be in the maintenance phase.

In DIV6 SN-L7MN co-cultures, we conducted quantitative live imaging of the bidirectional transport of lysosomal vesicles in SNs (Figure 1A-B). Briefly, confirming the formation of functional synapses in SN-L7MN cultures on DIV6, we incubated the cultures with LysoTracker Deep Red to stain lysosomal vesicles. LysoTracker Deep Red stains acidic organelles and is suitable for live imaging (Gonzalez-Polo, et al., 2005; Raben, et al., 2009). Kymograph analyses (Figure 1C) of transport measurements yielded the flux (J) (particles/µm/min) of transport, which assesses the number of particles that pass through a line across the axon over time, and the velocity (V) (µm/s) of transported organelles, which are related by the equation J=V*C (Badal et al., 2022).

**Figure 1.**
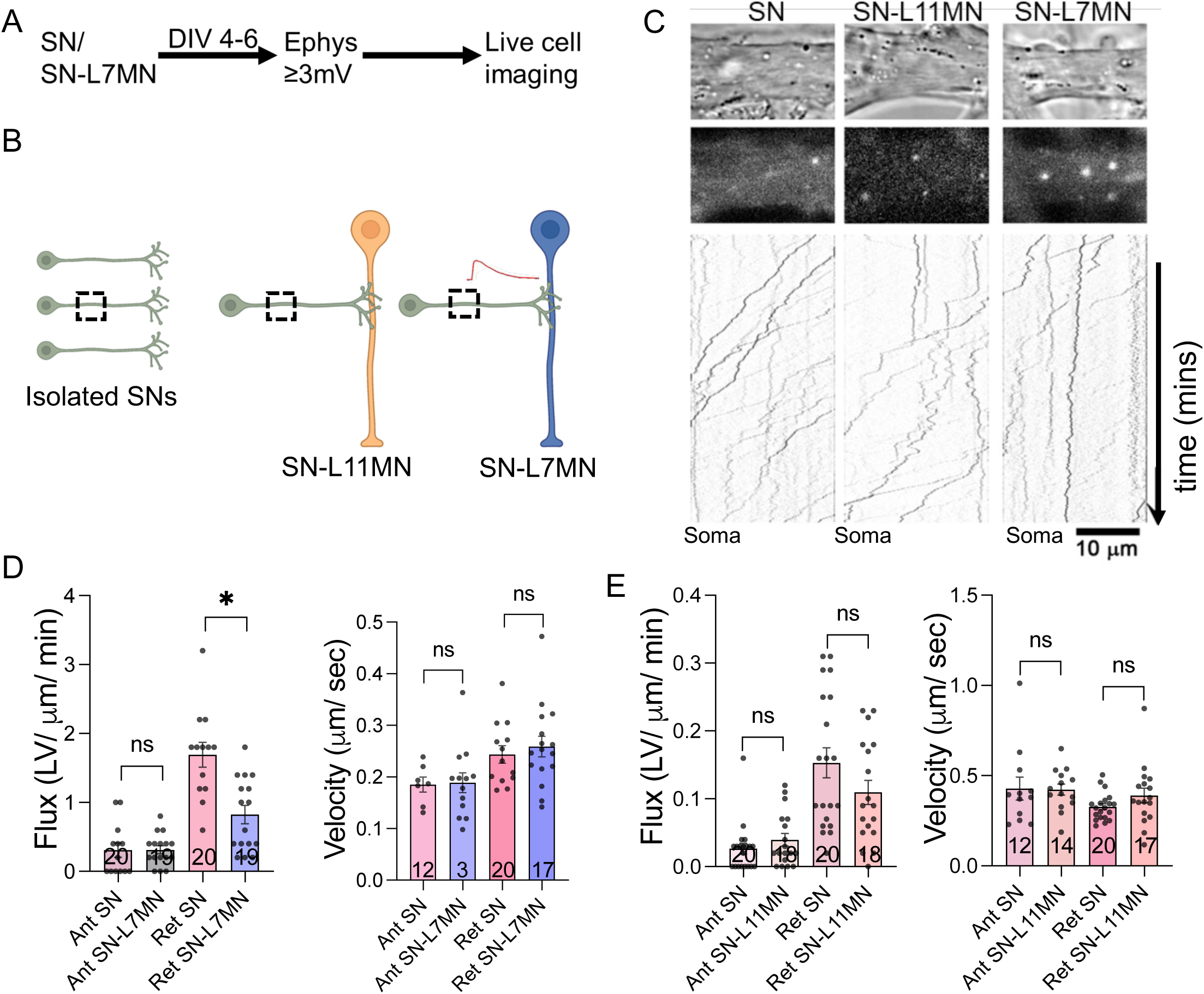
Synapse maintenance produces a decrease in the long-distance bidirectional flux of transport of lysosomal vesicles. **A.** Experimental strategy for live imaging of organelle transport from SN and SNL7MN cultures after SNL7MN electrophysiology during synapse maintenance. **B.** Cartoon schematic live transport imaging in SN, sensory neuron; L11MN, L11 motor neuron; SNL11MN, SN co-cultured with L11MN; L7MN, L7 motor neuron; SN-L7MN, SN co-cultured with L7MN during synapse maintenance. **C.** Confocal DIC and grayscale of fluorescent LROs particles within SN axons are shown. The fluorescence images in boxes show examples of regions where transport was analyzed. Representative kymographs of time-lapse axonal LV transport. Scale bar, 10um. **D.** Bar graphs show the flux and velocity of anterograde and retrograde LV transport in SNs and SNL7MNs analyzed by kymograph. The number of neurons analyzed in the experiment is indicated in the bar graphs. Error bars show SEMs. NS, nonsignificant; *p < 0.05. Student’s unpaired two-tailed t-test. **E.** Bars graph shows the flux and velocity of anterograde (Ant) and retrograde (Ret) LV transport in SNs and SNL11MN measured by kymograph analysis. The numbers of neurons analyzed in the experiment are indicated in the bar graphs. Error bars show SEMs. NS, nonsignificant. Student’s unpaired two-tailed t-test. Also see supplementary table S1.

SN-L7MN kymograph analyses revealed that lysosomal vesicle flux is reduced during basal synapse maintenance compared to transport in isolated SNs (Flux: SN retrograde lysosomal vesicle = 1.26 ± 0.17 SEM, n = 20 neurons, SN-L7MN retrograde lysosomal vesicle = 0.69 ± 0.15 SEM, n = 19 neurons, unpaired two-tailed t-test, p-value = 0.022; Velocity: SN retrograde lysosomal vesicle = 0.33 ± 0.02 SEM, n = 20 neurons, SN-L7MN retrograde lysosomal vesicle = 0.26 ± 0.03 SEM, n = 17 neurons, unpaired two-tailed t-test, p-value = 0.574; Figure 1C and 1D; Table S1; Video S1). These results suggest that synapse maintenance results in a significant and persistent reduction in the number of retrograde lysosomal vesicles in motion.

To determine whether the reduction in retrograde transport of lysosomal vesicles is indeed due to a functional synapse, we co-cultured SNs with L11MN, a non-target motor neuron with which SNs do not form a functional synapse. Quantitative analysis of transport in SNs versus SN-L11MN during DIV4-6 revealed that lysosomal vesicle flux and velocity were not significantly different (Flux: SN retrograde lysosomal vesicle = 0.15 ± 0.02 SEM, n = 20 neurons, SN-L11MN retrograde lysosomal vesicle = 0.11 ± 0.02 SEM, n = 18 neurons, unpaired two-tailed t-test, p-value = 0.237; Velocity: SN retrograde lysosomal vesicle = 0.33 ± 0.02 SEM, n = 20 neurons, SN-L11MN retrograde lysosomal vesicle = 0.39 ± 0.04 SEM, n = 17 neurons, unpaired two-tailed t-test, p-value = 0.15; Figure 1B, 1C, and 1E; Table S1; Video S1). Taken together, these results suggest that the formation of functional synapses between SN-L7MN results in a direction-specific reduction in the flux of lysosomal vesicle transport in SNs during synapse maintenance.

### Retrograde lysosomal vesicle flux is modulated during synapse maturation but not immediately after synapse formation

The modulation of lysosomal vesicle transport during synapse maintenance prompted us to investigate when this modulation begins following synapse formation. Therefore, we quantified the bidirectional transport of lysosomal vesicles at three time points: 6, 12, and 24 hours in vitro after synapse formation (Figures 2A and 2B). We found that lysosomal vesicle transport is not significantly modulated within 6 or 12 hours of synapse formation (6-hour flux: SN retrograde lysosomal vesicle = 0.32 ± 0.049 SEM, n = 18 neurons; SN-L7MN retrograde lysosomal vesicle = 0.266 ± 0.033 SEM, n = 39 neurons; unpaired two-tailed t-test, p-value = 0.36. Velocity: SN retrograde lysosomal vesicle = 0.439 ± 0.039 SEM, n = 18 neurons; SN-L7MN retrograde lysosomal vesicle = 0.434 ± 0.026 SEM, n = 35 neurons; unpaired two-tailed t-test, p-value = 0.91. 12-hour flux: SN retrograde lysosomal vesicle = 0.253 ± 0.038 SEM, n = 19 neurons; SN-L7MN retrograde lysosomal vesicle = 0.326 ± 0.045 SEM, n = 20 neurons; unpaired two-tailed t-test, p-value = 0.23. Velocity: SN retrograde lysosomal vesicle = 0.648 ± 0.07 SEM, n = 18 neurons; SN-L7MN retrograde lysosomal vesicle = 0.516 ± 0.05 SEM, n = 20 neurons; unpaired two-tailed t-test, p-value = 0.135; Figure 2C-F; Table S2, Video S2).

**Figure 2.**
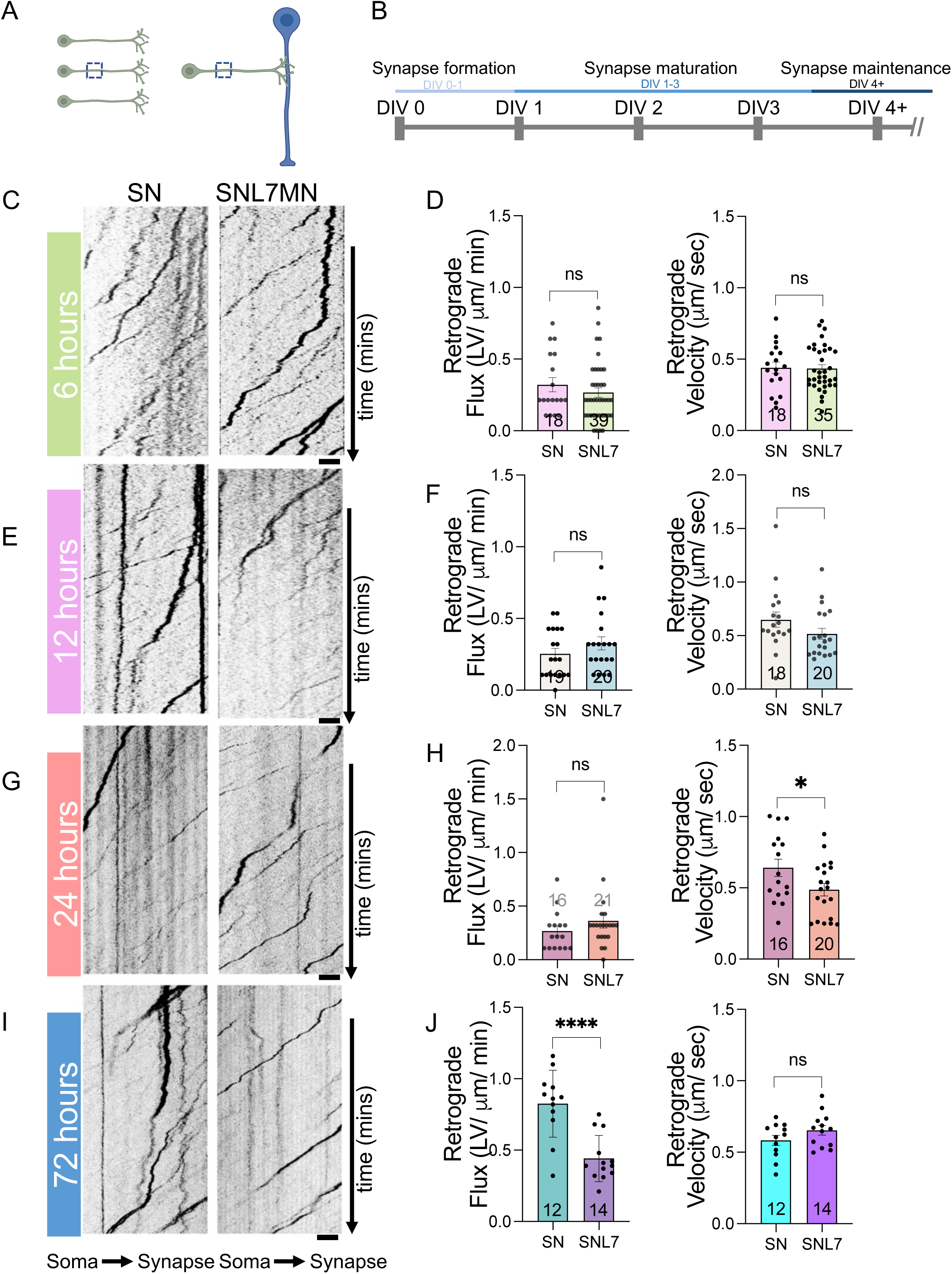
Dynamic changes in retrograde lysosomal vesicle transport during synapse formation and maturation. **A.** Cartoon schematic for imaging live LV transport in SN and SNL7MN during synapse formation and maturation. **B.** Timeline for SNL7MN *in vitro* cell-to-cell communication development. **C, E, G, I.** Representative kymographs of time-lapse axonal LROs transport at 6 **(C)**, 12 **(E)**, 24 **(G)**, and 72 **(I)** hours, respectively. Scale bar, 10um. **D, F, H, J**. Bar graphs show the flux and velocity of anterograde and retrograde LV transport in SN and SNL7MN at 6 **(D)**, 12 **(F)**, 24 **(H)**, and 72 **(J)** hours after plating analyzed by kymograph, respectively. The number of neurons analyzed in the experiment is indicated in the bar graphs. Error bars show SEMs. NS, nonsignificant; *p < 0.05. Student’s unpaired two-tailed t-test. Also see supplementary table S2.

At 24 hours post-synapse formation, lysosomal vesicle retrograde flux was not significantly modulated, but retrograde transport velocity showed a significant decrease (24-hour flux: SN retrograde lysosomal vesicle = 0.267 ± 0.045 SEM, n = 16 neurons; SN-L7MN retrograde lysosomal vesicle = 0.362 ± 0.066 SEM, n = 21 neurons; unpaired two-tailed t-test, p-value = 0.28. Velocity: SN retrograde lysosomal vesicle = 0.64 ± 0.06 SEM, n = 16 neurons; SN-L7MN retrograde lysosomal vesicle = 0.487 ± 0.04 SEM, n = 20 neurons; unpaired two-tailed t-test, p-value = 0.04; Figure 2G-2H; Table S2; Video S2).

Interestingly, our analysis of mitochondrial transport in SNs in these co-cultures showed that the flux of anterograde lysosomal vesicle transport became significantly enhanced at 24 hours. The flux of mitochondrial transport increased significantly in both anterograde and retrograde directions without significant changes in transport velocities. These observations suggest synapse formation results in organelle-specific and direction-specific modulation of long-distance transport.

Given that EPSPs stabilize within 72 hours of synapse formation, we next imaged lysosomal vesicle transport 72 hours after plating (Figure 2B). Kymograph transport analyses revealed that lysosomal vesicle retrograde flux is significantly reduced (72-hour flux: SN retrograde lysosomal vesicle = 0.824 ± 0.06, n = 12; SN-L7MN retrograde lysosomal vesicle = 0.441 ± 0.044, n = 14; unpaired two-tailed t-test, p-value = 0.00008. Velocity: SN retrograde lysosomal vesicle = 0.583 ± 0.03, n = 12; SN-L7MN retrograde lysosomal vesicle = 0.653 ± 0.03, n = 14; unpaired two-tailed t-test, p-value = 0.163; Figure 2I-2J; Table S2; Video S2). These results suggest that the modulation of lysosomal vesicle transport begins within 72 hours of synapse formation. Interestingly, the velocity changes observed at 24 hours became nonsignificant at 72 hours. Taken together, these results demonstrate a temporal and organelle-specific modulation of transport following synapse formation.

### Differential modulation of lysosomal vesicle and mitochondrial transport during long-term facilitation of SN-L7MN synapses

Since basal synapse formation and maintenance differentially modulate lysosomal vesicle transport (Figure 1-2), we investigated whether the transport of lysosomal vesicles is modulated by learning-associated formation of new synapses and remodeling of existing ones. The SN-L7MN synapses undergo long-term facilitation (LTF) in response to serotonin (5HT), a modulatory neurotransmitter involved in learning in Aplysia (Lin et al., 1994). In the intact animal, repeated tail shocks induce long-term sensitization (LTS), a form of non-associative learning, which results in the release of 5HT by serotonergic neurons to SN-L7MN synapses, leading to the formation of new synapses (Sutton et al., 2002; Wainwright et al., 2002). Importantly, exposure to 5HT in vitro also results in long-term enhancements in synaptic transmission (LTF) of SN-L7MN synapses and synaptic remodeling.

Therefore, we induced LTF in the SN-L7MN cultures by applying five interspaced pulses of serotonin (5×5HT) and simultaneously investigated the bidirectional transport of lysosomal vesicles at 1 and 24 hours after 5HT exposure. Transport of mitochondria was also quantified for comparison. Prior to imaging investigations, we performed electrophysiological experiments to verify LTF-associated enhancements in EPSPs (Figure 3A). Twenty four hours after 5xHT application EPSPs were significantly enhanced, substantiating 5×5HT leads to LTF in the SNL7MN system (EPSPs: Baseline EPSPs average = 11.1 ± 1.04 SEM, n = 21 neurons; post-5×5HT EPSPs average = 22.71 ± 1.88 SEM, n = 32 neurons; paired t-test, p = 0.0009; Figure 3B-3C; Table S3, Video S3).

**Figure 3.**
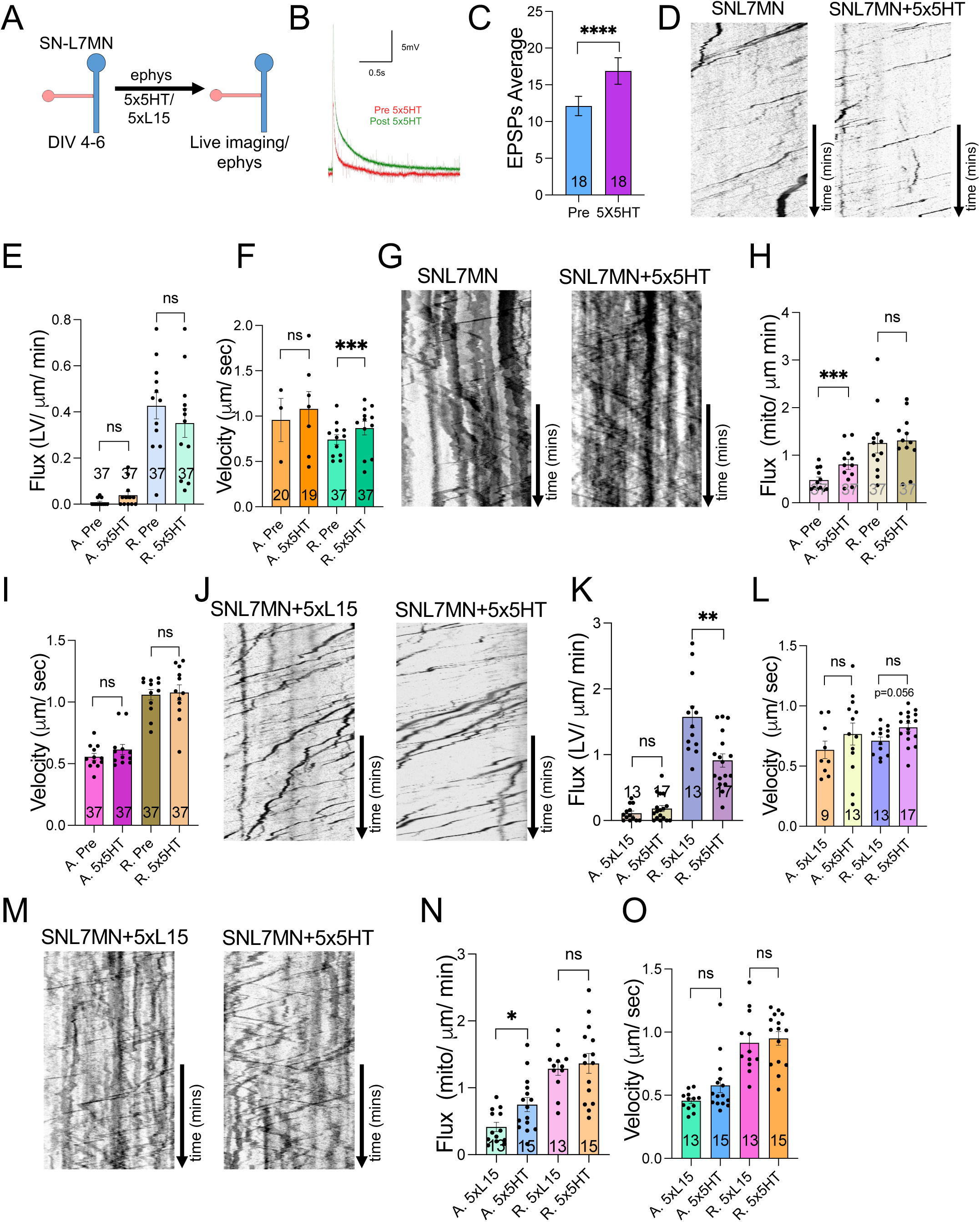
LTF Temporally and differentially regulates lysosomal vesicle and mitochondrial Transport. **A.** Experimental design for electrophysiological recordings before and 24 hours after 5×5HT. **B.** Representative trace of SNL7MN (DIV 4-6) excitatory postsynaptic potentials (EPSPs) before (green) and after (red) 5×5HT in millivolts (mV); scale display 5mV along the y-axis and 0.5 seconds (s) in the x-axis. **C.** Bar graph shows average excitatory postsynaptic potentials (EPSPs) in millivolts (mV) of SNL7MN before (pre) and 24 hours after 5×5HT application. The number of neurons analyzed in the experiment is indicated in the bar graphs. Error bars show SEMs. ****p < 0.0001. Student’s paired t-test. **D, G.** Representative kymographs of time-lapse axonal (**D)** LV and **(G)** mitochondrial transport. **E, F; H, I.** Bar graphs show anterograde and retrograde LV flux **(E)** and velocity **(F)**, and mitochondrial anterograde and retrograde flux **(H)** and velocity **(I)** before and 60 minutes after long-term facilitation induction (5×5HT) analyzed by kymograph. The number of neurons analyzed in the experiment is indicated in the bar graphs. Error bars show SEMs. NS, nonsignificant; ***p < 0.001. Student’s paired two-tailed t-test (flux). Student’s unpaired two-tailed t-test (velocity). **J, M.** Representative kymographs of time-lapse axonal LV **(J)** and mitochondrial **(M)** transport 24 hours after 5xL15 (control) or 5×5HT. **K, L, N, O.** Bar graphs show flux **(K)** and velocity **(L)** of anterograde and retrograde LV and flux **(N)** and velocity **(O)** of anterograde and retrograde mitochondrial transport 24 hours after 5xL15 (control) or 5×5HT application analyzed by kymograph. The number of neurons analyzed in the experiment is indicated in the bar graphs. Student’s unpaired two-tailed t-test. Error bars show SEMs. NS, nonsignificant; *p < 0.05, **p < 0.01.

Within 1 hour after 5×5HT, retrograde lysosomal vesicle velocity is enhanced, and anterograde mitochondrial flux is increased (60mins LTF: Flux: SN-L7MN retrograde lysosomal vesicle = 1.2 ± 0.08 SEM, n = 37 neurons; SN-L7MN+5×5HT retrograde lysosomal vesicle = 1.2 ± 0.07 SEM, n = 37 neurons; paired t-test, p-value = 0.925. Velocity: SN-L7MN retrograde lysosomal vesicle = 0.733 ± 0.019 SEM, n = 37 neurons; SN-L7MN+5×5HT retrograde lysosomal vesicle = 0.858 ± 0.029 SEM, n = 37 neurons; paired t-test, p-value = 0.00002. Mitochondria: Flux: SN-L7MN anterograde Mito = 0.451 ± 0.04 SEM, n = 37 neurons; SN-L7MN+5×5HT anterograde Mito = 0.736 ± 0.05 SEM, n = 37 neurons; paired t-test, p-value = 0.0003. SN-L7MN retrograde Mito = 1.01 ± 0.37 SEM, n = 37 neurons; SN-L7MN+5×5HT retrograde Mito = 1.14 ± 0.08 SEM, n = 37 neurons; paired t-test, p-value = 0.181. Velocity: SN-L7MN anterograde Mito = 0.456 ± 0.018 SEM, n = 37 neurons; SN-L7MN+5×5HT anterograde Mito = 0.521 ± 0.02 SEM, n = 37 neurons; paired t-test, p-value = 0.1267. SN-L7MN retrograde Mito = 0.902 ± 0.02 SEM, n = 37 neurons; SN-L7MN+5×5HT retrograde Mito = 1.06 ± 0.03 SEM, n = 37 neurons; paired t-test, p-value = 0.6139; Figure 3D-3I; Table S3; Video S3). These observations show dynamic, direction-specific modulations of retrograde lysosomal vesicle and anterograde mitochondrial transport during learning-associated synapse formation.

Next, we quantified transport 24 hours after LTF induction. At 24 hours post-5×5HT, lysosomal vesicle retrograde flux is further reduced compared to the vehicle (24hrs LTF: Flux: SN-L7MN+5XL15 retrograde lysosomal vesicle = 1.57 ± 0.16 SEM, n = 13 neurons; SN-L7MN+5×5HT retrograde lysosomal vesicle = 0.91 ± 0.101 SEM, n = 17 neurons; unpaired two-tailed t-test, p-value = 0.001. Velocity: SN-L7MN+5XL15 retrograde lysosomal vesicle = 0.71 ± 0.03 SEM, n = 13 neurons; SN-L7MN+5×5HT retrograde lysosomal vesicle = 0.82 ± 0.03 SEM, n = 17 neurons; unpaired two-tailed t-test, p-value = 0.02; unpaired two-tailed t-test, p-value = 0.0007; Figure 3J-3L; Table S3; Video S3). Only mitochondrial anterograde flux is regulated 24 hours after LTF (24hr LTF: Flux: SN-L7MN+5XL15 anterograde Mito = 0.414 ± 0.06 SEM, n = 13 neurons; SN-L7MN+5×5HT anterograde Mito = 0.746 ± 0.105 SEM, n = 15 neurons; unpaired two-tailed t-test, p-value = 0.01. SN-L7MN+5XL15 retrograde Mito = 1.29 ± 0.09 SEM, n = 13 neurons; SN-L7MN+5×5HT retrograde Mito = 1.36 ± 0.14 SEM, n = 15 neurons; unpaired two-tailed t-test, p-value = 0.706. Velocity: SN-L7MN+5XL15 anterograde Mito = 0.456 ± 0.02 SEM, n = 12 neurons; SN-L7MN+5×5HT anterograde Mito = 0.578 ± 0.05 SEM, n = 15 neurons; unpaired two-tailed t-test, p-value = 0.07. SN-L7MN+5XL15 retrograde Mito = 0.914 ± 0.06 SEM, n = 12 neurons; SN-L7MN+5×5HT retrograde Mito = 0.95 ± 0.05 SEM, n = 15 neurons; unpaired two-tailed t-test, p-value = 0.68; Figure 3M-3O; Table S3; Video S3). These data demonstrate that learning-associated synapse formation temporally and persistently regulates retrograde lysosomal vesicle and anterograde mitochondrial trafficking. Together, these results establish that bidirectional long-distance transport is temporally modulated in an organelle-specific and direction-specific manner during synapse formation, maintenance, and plasticity.

### Assessing the role of the postsynaptic neuron in regulating presynaptic lysosomal vesicle transport during synapse maintenance

In our search for mechanisms underlying the reduction in long-distance retrograde transport of lysosomal vesicles during synapse maintenance, we first considered the possibility that persistent signals from the postsynaptic neuron (L7MN) modulate presynaptic transport (Figure 4A). To investigate the contribution of L7MN cell body-generated signals in regulating presynaptic lysosomal vesicle transport, we microdissected the L7MN during synapse maintenance and then imaged presynaptic axonal transport. An increase in presynaptic lysosomal vesicle transport after L7MN cell body removal (CBR) would suggest a role for L7MN cell body-generated signaling in modulating presynaptic transport. Therefore, we microdissected the L7MN cell body and measured lysosomal vesicle transport 24 hours later (Figure 4B). Quantitative transport analyses indicated no significant changes in lysosomal vesicle flux or velocity in SNs with L7MN-CBR compared to SNs with an intact postsynaptic L7MN (24hr CBR: SN retrograde lysosomal vesicle = 1.68 ± 0.26 SEM, n = 11 neurons, SNL7MN retrograde lysosomal vesicle = 1.45 ± 0.26 SEM, n = 10 neurons, unpaired two-tailed t-test, p-value = 0.69; Velocity: SN retrograde lysosomal vesicle = 0.63 ± 0.06 SEM, n = 8 neurons, SNL7MN retrograde lysosomal vesicle = 0.66 ± 0.04 SEM, n = 10 neurons, unpaired two-tailed t-test, p-value = 0.54; Figure 4B-4C; Table S4). These results suggest that postsynaptic cell body-generated signals do not regulate presynaptic long-distance transport of Lysosomal vesicles during synapse maintenance.

**Figure 4.**
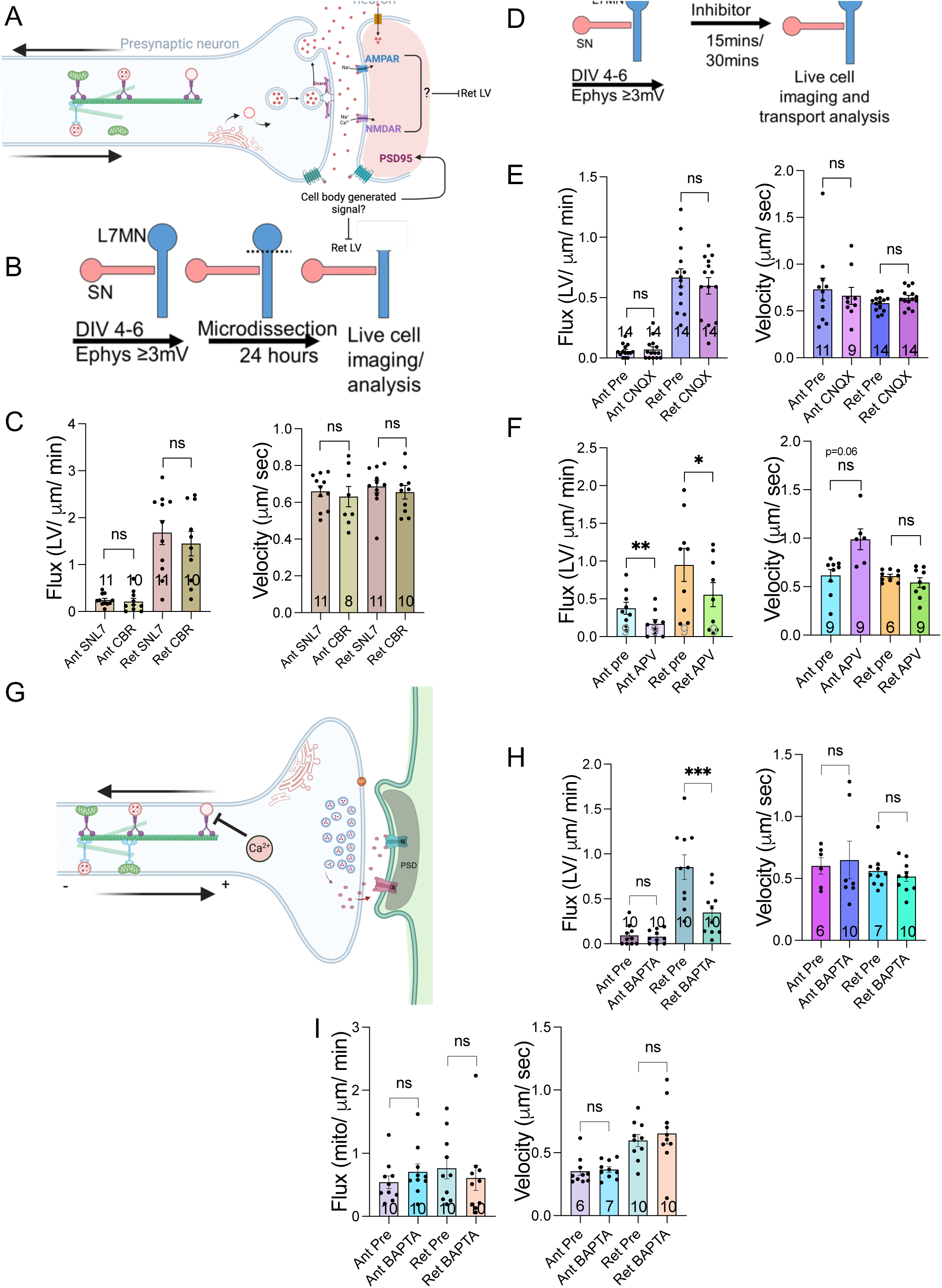
Role of the postsynaptic neuron in modulating presynaptic transport of lysosomal vesicles during synapse maintenance. **A.** Cartoon of experimental design and predicted mechanism that reduced retrograde LRO transport. Mechanism 1 predicts that L7 postsynaptic cell body-generated signals regulate presynaptic retrograde (ret) LV. Mechanism 2 predicts that AMPA and NMDA receptor signaling regulates presynaptic retrograde (ret) LV transport **B.** Experimental design for postsynaptic soma removal experiments. **C.** Bar graphs show the flux and velocity of anterograde and retrograde LV transport with intact MN (CB) or with MN cell body removed (CBR) analyzed by kymograph. The numbers of neurons analyzed in the experiment are indicated in the bar graphs. Error bars show SEMs. NS, nonsignificant. Student’s unpaired two-tailed t-test. **D.** Experiment design for inhibiting glutamatergic receptor signaling. **E, F.** Bar graphs show the flux and velocity of anterograde and retrograde LROs transport before and after CNQX **(E)** or APV **(F)** treatment analyzed from kymographs. The number of neurons analyzed in the experiment is indicated in the bar graphs. Error bars show SEMs. NS, nonsignificant; **p < 0.01. Student’s paired t-test (CNQX Flux, APV Flux); Student’s unpaired two-tailed t-test (CNQX Velocity and APV Velocity). **G.** Cartoon of experimental design and predicted mechanism that Ca^2+^ signaling regulates retrograde LV transport. **H, I.** Bar graphs show flux and velocity of anterograde and retrograde LV **(H)** and mitochondrial **(I)** transport before and after BAPTA application analyzed from kymographs. The number of neurons analyzed in the experiment is indicated in the bar graphs. Student’s paired t-test. Error bars show SEMs. NS, nonsignificant; **p < 0.01, ***p < 0.001.

Next, we examined the role of signaling from two important postsynaptic glutamatergic receptors, AMPA (α-amino-3-hydroxy-5-methyl-4-isoxazolepropionic acid) and NMDA (N-methyl-D-aspartate), in regulating presynaptic lysosomal vesicle retrograde transport. First, we utilized cyanquixaline (CNQX), an AMPAR antagonist that reduces AMPAR-mediated synaptic transmission, in SNL7MN to investigate whether AMPAR signaling modulates presynaptic axonal lysosomal vesicle transport (Figure 4A, 4D). Quantitative transport analyses 30 minutes after CNQX bath application demonstrated no significant changes in lysosomal vesicle flux or velocity when comparing transport in SNL7MNs before and after CNQX application (CNQX: pre-CNQX SNL7MN retrograde lysosomal vesicle = 0.66 ± 0.07 SEM, n = 14 neurons, post-CNQX SNL7MN retrograde lysosomal vesicle = 0.598 ± 0.06 SEM, n = 14 neurons, paired t-test, p-value = 0.44; Velocity: pre-CNQX SNL7MN retrograde lysosomal vesicle = 0.583 ± 0.02 SEM, n = 14 neurons, post-CNQX SNL7MN retrograde lysosomal vesicle = 0.639 ± 0.02 SEM, n = 14 neurons, paired t-test, p-value = 0.14; Figure 4E; Table S4). These data suggest that the immediate inhibition of synaptic transmission via CNQX inhibition of glutamatergic AMPA receptors does not affect the presynaptic transport of lysosomal vesicles.

Subsequently, we studied the effects of the NMDA receptor antagonist (DL)-2-amino-5-phosphonovaleric acid (APV) on lysosomal vesicle transport in SNL7MN for 15 minutes (Figure 4A, 4D). Quantitative transport analyses revealed a significant reduction in lysosomal vesicle flux but no changes in velocity when comparing transport in SNs before and after APV application (APV: pre-APV SNL7MN anterograde lysosomal vesicle = 0.375 ± 0.07 SEM, n = 9 neurons, post-APV SNL7MN anterograde lysosomal vesicle = 0.168 ± 0.05 SEM, n = 9 neurons, paired t-test, p-value = 0.006; pre-APV SNL7MN retrograde lysosomal vesicle = 0.948 ± 0.219 SEM, n = 9 neurons, post-APV SNL7MN retrograde lysosomal vesicle = 0.5539 ± 0.159 SEM, n = 9 neurons, paired t-test, p-value = 0.02; Velocity: pre-APV SNL7MN retrograde lysosomal vesicle = 0.608 ± 0.01 SEM, n = 9 neurons, post-APV SNL7MN retrograde lysosomal vesicle = 0.542 ± 0.05 SEM, n = 9 neurons, paired t-test, p-value = 0.143; Figure 4F; Table S4). While we observed changes in presynaptic lysosomal vesicle transport with APV-inhibited synaptic transmission, we suspect these results are due to secondary effects of the pharmacological reagent. For example, others observed secondary effects, such as reduced Ca2+ signaling at presynaptic sites. Therefore, we next investigated whether Ca2+ signaling regulates lysosomal vesicle axonal transport during synapse maintenance (Figure 4G). We used a calcium chelator, BAPTA AM [1mM], for 30 minutes to reduce intracellular Ca2+ levels in SNL7MN and observed lysosomal vesicle transport. lysosomal vesicle retrograde flux was significantly reduced with BAPTA application, however, we found mitochondrial transport was not affected (BAPTA: pre-BAPTA SNL7MN retrograde lysosomal vesicle = 0.853 ± 0.136 SEM, n = 10 neurons, post-BAPTA SNL7MN retrograde lysosomal vesicle = 0.346 ± 0.07 SEM, n = 10 neurons, paired t-test, p-value = 0.0008; Velocity: pre-BAPTA SNL7MN retrograde lysosomal vesicle = 0.559 ± 0.04 SEM, n = 10 neurons, post-BAPTA SNL7MN retrograde lysosomal vesicle = 0.514 ± 0.039 SEM, n = 10 neurons, paired t-test, p-value = 0.309; Figure 4H-I; Table S4). Thus, during synapse maintenance, lysosomal vesicle transport is regulated by intracellular Ca2+ signaling.

### The role of plasticity-related signaling pathways in the modulation of transport of lysosomal vesicles

Since our quantification of lysosomal vesicle transport did not show a role for NMDAR, AMPAR and L7MN cell body-generated signals during synapse maintenance, we concluded that synapse formation results in persistent modification of lysosomal vesicle transport during synapse maintenance. We assumed that specific intracellular signaling pathways might modulate persistent reduction in retrograde lysosomal vesicle flux following synapse formation. Quantitative analyses of transport during formation, maintenance, and plasticity suggest a correlation with synapse maintenance with reduction in lysosomal vesicle transport. This could be due to the increase in the metabolic demand for maintaining synaptic connections. In order to meet this demand, neurons must optimally use resources by re-allocation. Alternately, fewer materials need to be transported back to the cell body because available resources are efficiently consumed at the synapse. Therefore, we assessed that signaling pathways involved in plasticity might modulate lysosomal vesicle transport.

Utilizing isolated SNs, we investigated the particular roles of signaling pathways: protein kinase C (PKC), intracellular calcium release from inositol triphosphate (IP3) receptors (IP3R) on the endoplasmic reticulum (ER), cyclic AMP (cAMP) and protein kinase A (PKA) signaling on SN axonal lysosomal vesicles transport. First, we expose isolated SNs to either PMA (phorbol12-myristate13-acetate, PKC signaling activator, concentration 50nM) for 15min, Adenophostin A (ADA, activator of IP3R signaling, concentration 10mM) for 15min, forskolin (FK) (cAMP signaling activator, concentration 50mM) for 30min, or 14-22 amide (PKAi) (PKA signaling inhibitor, concentration 360nM) for 30min (Figure S2A-S2B). This was followed by analyses of lysosomal vesicle transport. If these signaling pathways regulate retrograde lysosomal vesicle transport, then we should observe a significant reduction in retrograde lysosomal vesicle transport after phrenological activation. However, we found PKC, ADA, FK activation, and PKAi did not affect lysosomal vesicle transport in isolated SNs (SNs: pre-PMA retrograde lysosomal vesicle flux = 0.381 ± 0.05 SEM, n = 16 neurons: 0.38, post-PMA = 0.38 ± 0.06 SEM, n = 17 neurons, paired t-test, p-value = 0.808; pre-ADA retrograde lysosomal vesicle flux = 0.361 ± 0.089 SEM, n = 17 neurons, post-ADA flux = 0.274 ± 0.067 SEM, n = 17 neurons, paired t-test, p-value = 0.354; pre-FK retrograde lysosomal vesicle flux = 0.367 ± 0.107 SEM, n = 6 neurons, post-FK flux = 0.328 ± 0.1 SEM, n = 6 neurons, paired t-test, p-value = 0.811; pre-PKAi retrograde lysosomal vesicle flux = 0.795 ± 0.08 SEM, n = 16 neurons, post-PKAi flux = 0.716 ± 0.08 SEM, n = 16 neurons, paired t-test, p-value = 0.533; Figure S2C-S2F; Table S10). Briefly, mitochondrial transport was not modulated with PKAi in SN (Figure S3G; Table S10). These results indicate that lysosomal vesicle transport is not regulated in SN by plasticity-related signaling pathways (PKC, IP3R, cAMP, PKA) in the absence of a functional synapse.

### Assessing the role of protein degradation and autophagy in modulating lysosomal vesicle transport

Given the key role lysosomes play in mediating protein degradation and autophagy, we considered whether a reduction in protein degradation or inhibition of autophagy during synapse maintenance might underlie the observed reduction in retrograde flux of lysosomal vesicle transport. To investigate this, we used inhibitors of protein degradation, mTOR signaling, and autophagy in our imaging experiments. These experiments were conducted in isolated SNs and SN-L7MN neurons, imaging the transport of lysosomal vesicles and mitochondria in DIV 4-6 neurons (Figure 5A-B).

**Figure 5.**
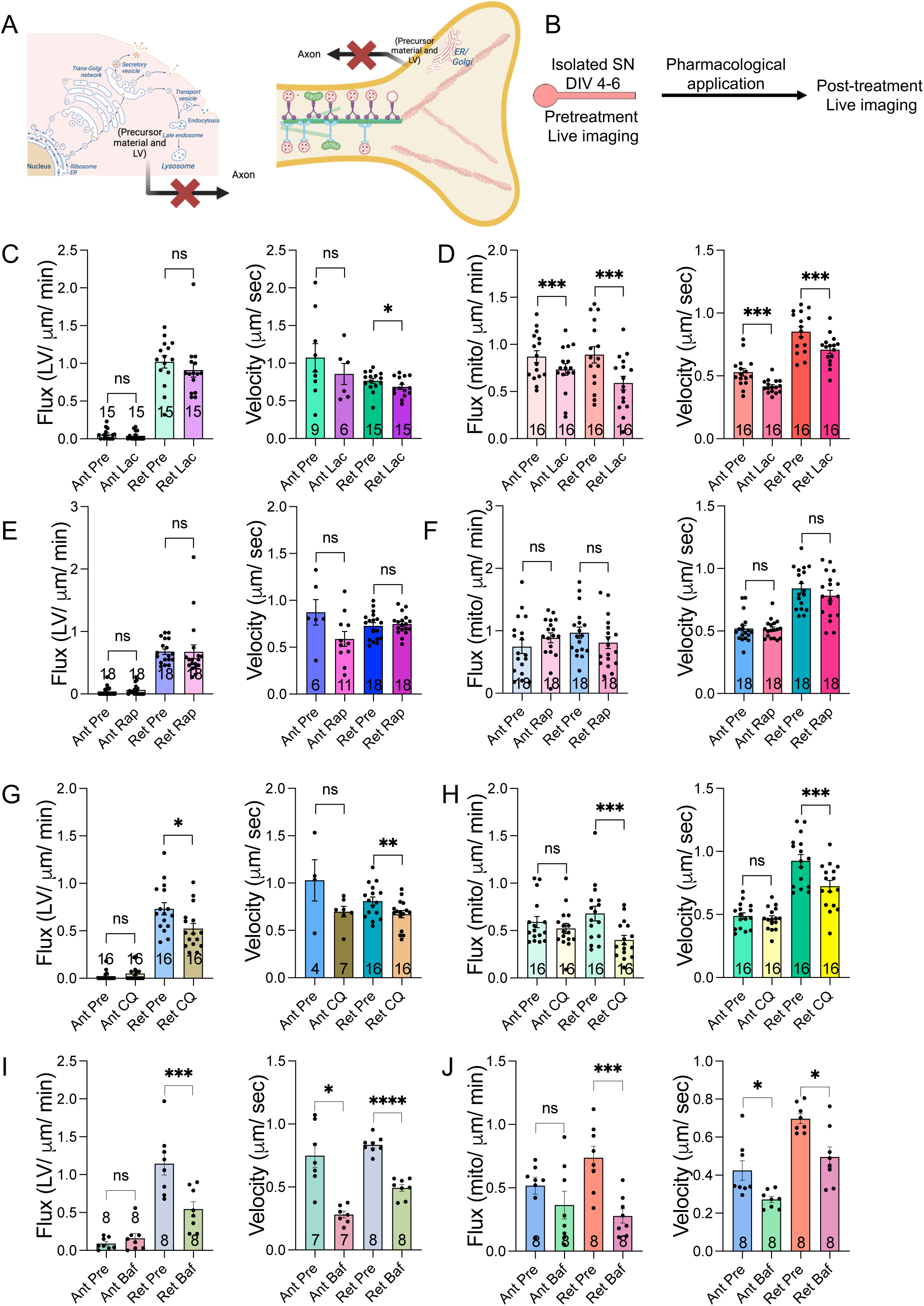
Effect of inhibition of protein degradation, mTOR signaling and autophagy on the bidirectional long distance transport in the absence of a synapse. **A.** Pharmacological interventions to assess signaling pathways modulating retrograde transport of lysosomal vesicles. **B.** Experiment design. **C-J.** Bar graphs show flux and velocity of anterograde and retrograde lysosomal vesicle **(C, E, G, I)** and mitochondrial **(D, F, H, J)** transport in SNs before and after Lactacystin (Lac) **(C, JD)**, Rapamycin (Rap) **(E, F)**, Chloroquine (CQ) **(G, H)** and Bafilomycin (Baf) **(I, J)** treatments analyzed by kymograph. The number of neurons analyzed in the experiment is indicated in the bar graphs. Error bars show SEMs. NS, nonsignificant; *p < 0.05, **p < 0.01, ***p < 0.001. Student’s unpaired two-tailed t-test.

Briefly, isolated SNs (DIV4-6) were treated with lactacystin (a proteasome and non-lysosomal intracellular protein degradation inhibitor), rapamycin (an mTOR signaling inhibitor), chloroquine (CQ, an autophagy inhibitor), and bafilomycin A (Baf, which inhibits autophagic flux and autophagosome-lysosome fusion). Rapamycin inhibits the mechanistic target of rapamycin (mTOR) signaling and promotes autophagy (Sekiguchi et al., 2012). Both CQ and Baf inhibit autophagolysosomal formation by preventing autophagosome fusion with lysosomes, although Baf does not alter Golgi stack or organization, despite disrupting the trans-Golgi network (TGN).

We imaged lysosomal vesicle and mitochondrial transport before and 30 minutes after treating neurons with 50 µM lactacystin. Lactacystin did not alter lysosomal vesicle flux, but it reduced retrograde lysosomal vesicle velocity. However, it reduced both the flux and velocity of bidirectional mitochondrial transport. Specifically, the retrograde lysosomal vesicle flux remained unchanged (pre-Lactacystin: 1.02 ± 0.08 SEM, n = 15 neurons; post-Lactacystin: 0.912 ± 0.09 SEM, n = 15 neurons; p = 0.318), while retrograde lysosomal vesicle velocity decreased (pre-Lactacystin: 0.766 ± 0.03 SEM, n = 15 neurons; post-Lactacystin: 0.686 ± 0.03 SEM, n = 15 neurons; p = 0.03; Figure 5C-D; Table S5). For mitochondrial flux, anterograde flux was reduced (pre-Lactacystin: 0.87 ± 0.06 SEM, n = 16 neurons; post-Lactacystin: 0.735 ± 0.06 SEM, n = 16 neurons; p = 0.009), as was retrograde flux (pre-Lactacystin: 0.888 ± 0.08 SEM, n = 16 neurons; post-Lactacystin: 0.59 ± 0.08 SEM, n = 16 neurons; p = 0.004). Mitochondrial velocity showed similar reductions (anterograde: pre-Lactacystin: 0.529 ± 0.02 SEM, n = 16 neurons; post-Lactacystin: 0.417 ± 0.01 SEM, n = 16 neurons; p = 0.002; retrograde: pre-Lactacystin: 0.851 ± 0.16 SEM, n = 16 neurons; post-Lactacystin: 0.7 ± 0.02 SEM, n = 16 neurons; p = 0.004; Figure 5C-D; Table S5). Thus, protein degradation inhibition impacts the bidirectional transport of mitochondria but not lysosomal vesicles. In contrast, inhibition of mTOR signaling with rapamycin did not affect the flux or velocity of lysosomal vesicle and mitochondrial transport, suggesting that brief inhibition of mTOR signaling does not impair their bidirectional transport (Figure 5E-5F).

Interestingly, CQ application reduced both retrograde lysosomal vesicle flux and velocity. Retrograde lysosomal vesicle flux decreased (pre-CQ: 0.730 ± 0.06 SEM, n = 16 neurons; post-CQ: 0.524 ± 0.05 SEM, n = 16 neurons; p = 0.025), as did retrograde lysosomal vesicle velocity (pre-CQ: 0.809 ± 0.04 SEM, n = 16 neurons; post-CQ: 0.67 ± 0.03 SEM, n = 16 neurons; p = 0.005; Figure 5G; Table S5). Similarly, retrograde mitochondrial transport was also reduced by CQ (flux: pre-CQ: 0.679 ± 0.07 SEM, n = 16 neurons; post-CQ: 0.399 ± 0.04 SEM, n = 16 neurons; p = 0.001; velocity: pre-CQ: 0.925 ± 0.04 SEM, n = 16 neurons; post-CQ: 0.724 ± 0.04 SEM, n = 16 neurons; p = 0.001; Figure 5H; Table S5). Thus, in isolated SNs, CQ disrupts retrograde transport of both mitochondria and lysosomal vesicles.

To further investigate the role of autophagosome fusion in reducing lysosomal vesicle transport, we applied Baf [2 µM] for 30 minutes to SNs and imaged lysosomal vesicle and mitochondrial transport. Baf significantly reduced lysosomal vesicle and mitochondrial retrograde flux and bidirectional velocity (SN Baf: Flux: pre-Baf SN retrograde lysosomal vesicle = 1.14 ± 0.14 SEM, n = 8 neurons; post-Baf SN retrograde lysosomal vesicle = 0.543 ± 0.09 SEM, n = 8 neurons; paired t-test, p = 0.008. Velocity: pre-Baf SN anterograde lysosomal vesicle = 0.748 ± 0.09 SEM, n = 7 neurons; post-Baf SN anterograde lysosomal vesicle = 0.28 ± 0.02 SEM, n = 7 neurons; paired t-test, p = 0.01. Pre-Baf SN retrograde lysosomal vesicle = 0.834 ± 0.02 SEM, n = 8 neurons; post-Baf SN retrograde lysosomal vesicle = 0.49 ± 0.02 SEM, n = 8 neurons; paired t-test, p = 4.11E-06. SN Baf: Flux: pre-Baf SN retrograde Mito = 0.73 ± 0.09 SEM, n = 8 neurons; post-Baf SN retrograde Mito = 0.276 ± 0.05 SEM, n = 8 neurons; paired t-test, p = 0.002. Velocity: pre-Baf SN anterograde Mito = 0.424 ± 0.05 SEM, n = 8 neurons; post-Baf SN anterograde Mito = 0.272 ± 0.01 SEM, n = 8 neurons; paired t-test, p = 0.02. Pre-Baf SN retrograde Mito = 0.696 ± 0.02 SEM, n = 8 neurons; post-Baf SN retrograde Mito = 0.495 ± 0.05 SEM, n = 8 neurons; paired t-test, p = 0.01; Figure 5I-J; Table S5). Therefore, inhibition of autophagy by Baf disrupts retrograde flux of mitochondria without interfering with their anterograde flux.

Next, we examined whether Lact, Rapa, CQ, and Baf produce similar effects on bidirectional transport in SNs during synapse maintenance. Initially, we applied Lact, Rapa, and CQ to SNL7MN and measured their effects on synaptic transmission (Figure 6A, Figure S3). Lact, Rapa, and CQ significantly reduced synaptic transmission (EPSPs: Pre-Lact EPSPs average = 16.44 ± 2.387 SEM, n = 9 neurons; post-Lact EPSPs average = 10.89 ± 1.889 SEM, n = 9 neurons; paired t-test, p = 0.0001. EPSPs: Pre-Rapa EPSPs average = 15.5 ± 2.96 SEM, n = 10 neurons; post-Rapa EPSPs average = 8.5 ± 1.16 SEM, n = 10 neurons; paired t-test, p = 0.02. EPSPs: Pre-CQ EPSPs average = 14.73 ± 2.17 SEM, n = 15 neurons; post-CQ EPSPs average = 3.7 ± 0.58 SEM, n = 15 neurons; unpaired t-test, p = 0.005; Figure 6A-C, Figure S3A-D; Table S6, Table S10), indicating the roles of autophagy, mTOR signaling, and autophagosome fusion in maintaining synaptic transmission.

**Figure 6.**
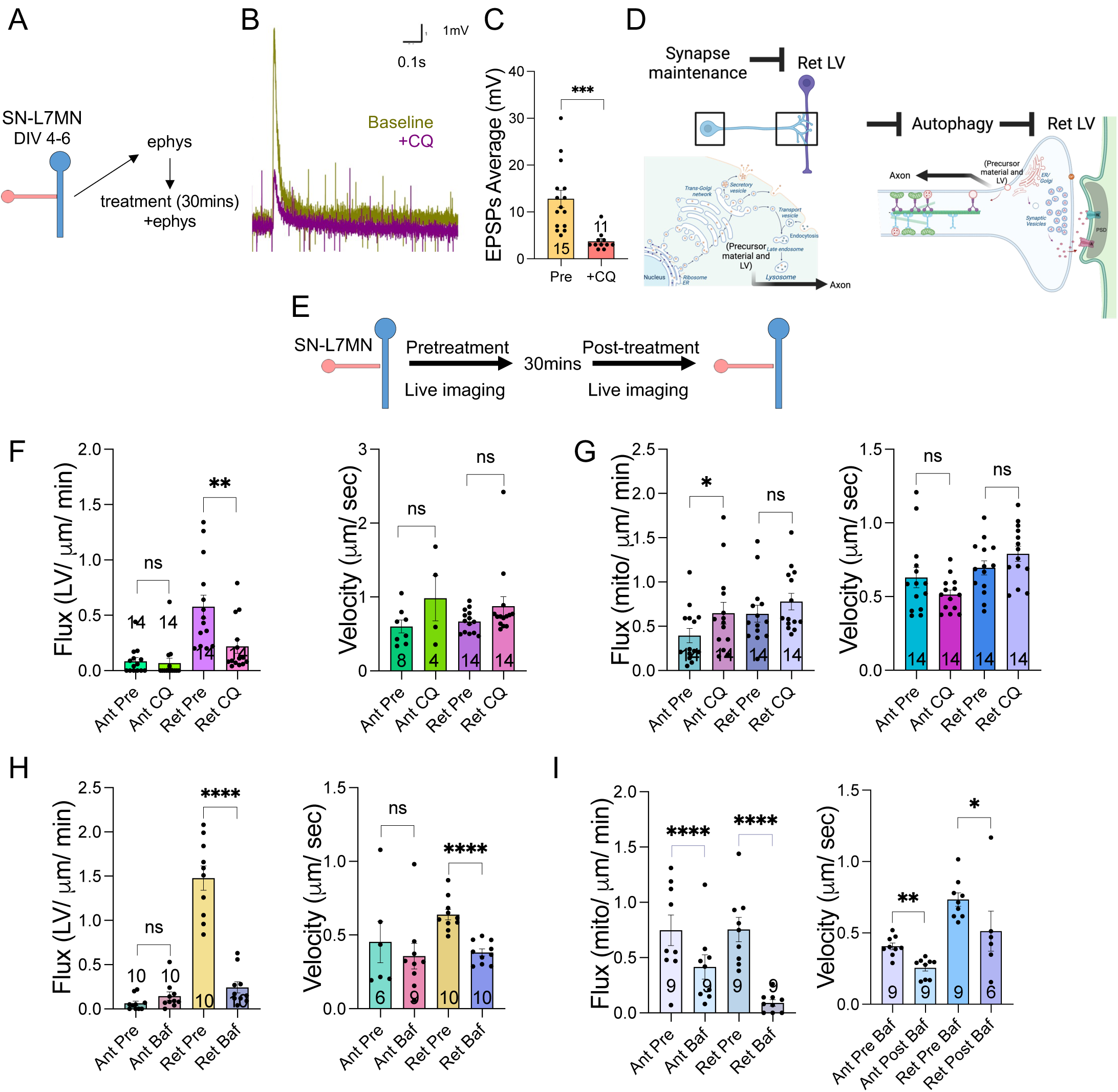
Transport of lysosomal vesicles is regulated by autophagy. **A.** Experiment design of electrophysiological recordings before and 30 minutes (mins) after pharmacological application. **B.** Representative trace of SNL7MN (DIV 4-6) excitatory postsynaptic potentials (EPSPs) in SNL7MN before (brown) and after (purple) chloroquine (+CQ), in millivolts (mV); scale display mV along the y-axis and 0.1 seconds (s) in the x-axis. **C.** Bar graph shows average EPSPs in mV of SNL7MN before (pre) and 30 mins +CQ. The number of neurons analyzed in the experiment is indicated in the bar graphs. Error bars show SEMs. **p < 0.01. ***p < 0.001. Student’s paired t-test. **D.** Cartoon of possible mechanism modulate retrograde LV transport. **E.** Experiment design. **F-I.** Bar graphs show the flux and velocity of anterograde and retrograde LV **(F, H)** and mitochondrial **(G, I)** transport in SNL7MNs before and after Chloroquine (CQ) **(F, G)** and bafilomycin **(H, I)** treatment analyzed by kymograph. The number of neurons analyzed in the experiment is indicated in the bar graphs. Error bars show SEMs. NS, nonsignificant; *p < 0.05, **p < 0.01, ***p < 0.001, ****p < 0.0001. Student’s unpaired two-tailed t-test.

Considering the significant reductions in synaptic communication following CQ application, we imaged lysosomal vesicle and mitochondrial transport in SNL7MN neurons treated with CQ (Figure 6D-E). Thirty minutes of CQ treatment in SNL7MN resulted in reduced lysosomal vesicle retrograde flux and enhanced mitochondrial anterograde flux (CQ lysosomal vesicle : Flux: pre-CQ SNL7MN retrograde lysosomal vesicle = 0.577 ± 0.105, n = 14; post-CQ SNL7MN retrograde lysosomal vesicle = 0.218 ± 0.06, n = 14; paired t-test, p = 0.01. CQ Mito: Flux: pre-CQ SNL7MN anterograde Mito = 0.39 ± 0.08, n = 14; post-CQ SNL7MN anterograde Mito = 0.65 ± 0.12, n = 14; paired t-test, p = 0.013. Pre-CQ SNL7MN retrograde Mito = 0.64 ± 0.1, n = 14; post-CQ SNL7MN retrograde Mito = 0.78 ± 0.09, n = 14; paired t-test, p = 0.08; Figure 6F-6G; Table S6; Video S4). These reductions in lysosomal vesicle trafficking indicate that the endo-lysosomal transport system regulates presynaptic lysosomal vesicle trafficking in SNs.

We followed up on these investigations with Baf application to SNL7MN neurons (Figure 6H-6I). Baf exposure also reduced retrograde lysosomal vesicle flux and velocity and decreased bidirectional mitochondrial flux and velocity (SNL7MN Baf: Flux: pre-Baf retrograde lysosomal vesicle = 1.47 ± 0.13 SEM, n = 10 neurons; post-Baf retrograde lysosomal vesicle = 0.242 ± 0.06 SEM, n = 10 neurons; paired t-test, p = 7.04E-05. Velocity: pre-Baf retrograde lysosomal vesicle = 0.638 ± 0.03 SEM, n = 10 neurons; post-Baf retrograde lysosomal vesicle = 0.381 ± 0.02 SEM, n = 10 neurons; paired t-test, p = 0.0002. Flux: pre-Baf anterograde Mito = 0.74 ± 0.13 SEM, n = 9 neurons; post-Baf anterograde Mito = 0.109 ± 0.1 SEM, n = 9 neurons; paired t-test, p = 0.004. Pre-Baf retrograde Mito = 0.75 ± 0.11 SEM, n = 9 neurons; post-Baf retrograde Mito = 0.09 ± 0.08 SEM, n = 9 neurons; paired t-test, p = 0.0003. Velocity: pre-Baf anterograde Mito = 0.406 ± 0.02 SEM, n = 9 neurons; post-Baf anterograde Mito = 0.256 ± 0.069 SEM, n = 9 neurons; paired t-test, p = 0.002. Pre-Baf retrograde Mito = 0.733 ± 0.04 SEM, n = 9 neurons; post-Baf retrograde Mito = 0.512 ± 0.14 SEM, n = 9 neurons; unpaired t-test, p = 0.05; Figure 6H-6I; Table S6; Video S4), indicating a key role for autophagy in modulating mitochondrial and lysosomal vesicle transport during synapse maintenance.

### Organelle-specific modulation of transport by endoplasmic reticulum (ER) to golgi transport and membrane-material availability

We continued our experiments to identify mechanisms that specifically modulate the transport of lysosomal vesicles. Since the ER-Golgi system is a hub for membrane protein sorting and is responsible for synthesizing trafficking vesicles, including precursors used in exocytosis and endocytosis pathways, we inhibited ER to Golgi transport of proteins using Brefeldin A (Figure 7A-8B). Interestingly, BFA application (60 minutes) enhanced the flux of anterograde mitochondrial transport without altering the flux or velocities of lysosomal vesicle transport (BFA lysosomal vesicle : Flux: pre-BFA SNL7MN retrograde lysosomal vesicle = 0.87 ± 0.07, n = 8; post-BFA SNL7MN retrograde lysosomal vesicle = 0.72 ± 0.09, n = 8; paired t-test, p = 0.14; Figure 7B-8C; Table S7).

**Figure 7.**
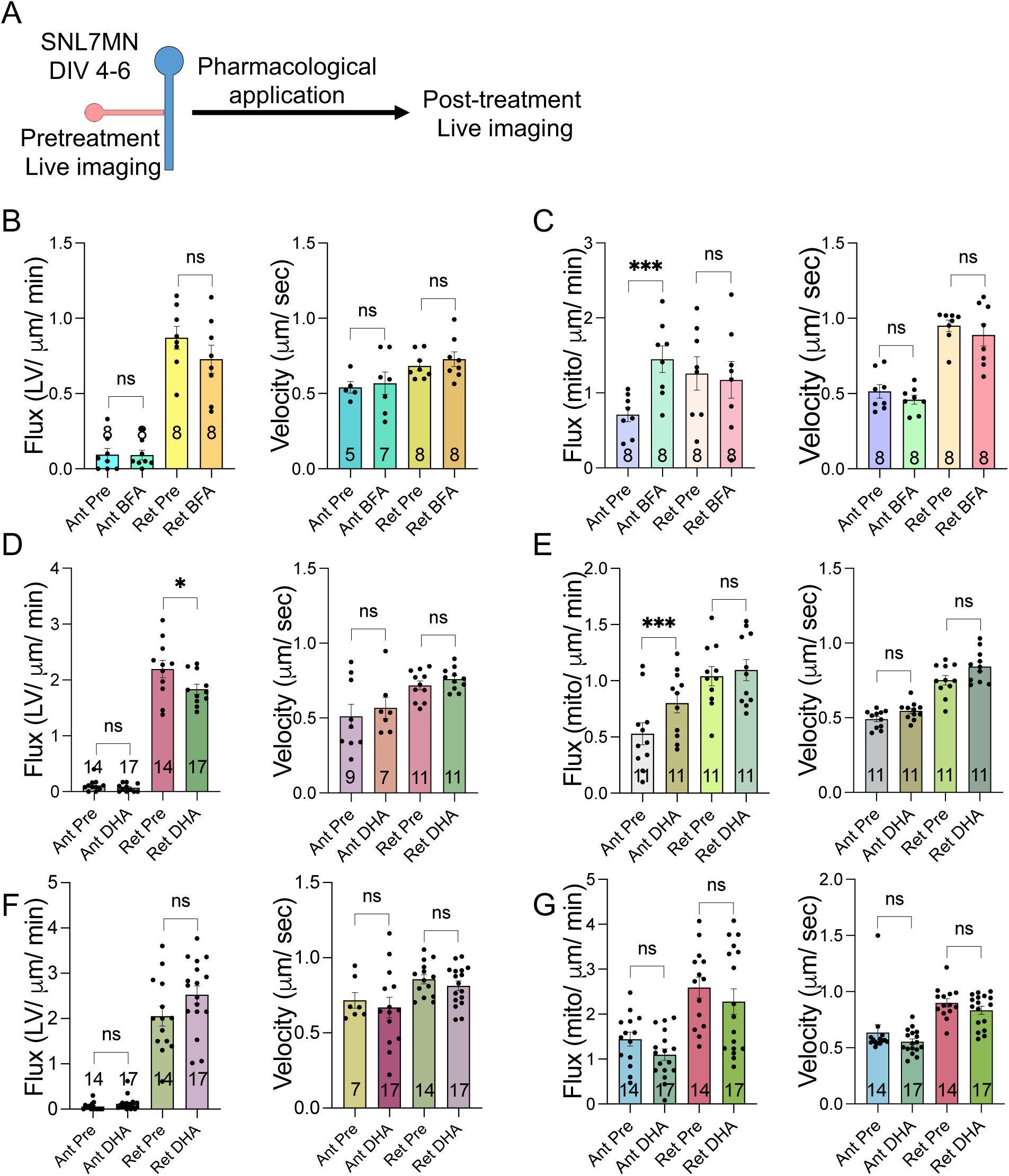
Assessing the role of ER-golgi transport and effect of DHA on long-distance bidirectional transport of mitochondria and lysosomal vesicles. **A.** Cartoon of experimental design. **B-C.** Bar graphs show the flux and velocity of anterograde and retrograde LV **(B)** and mitochondrial **(C)** transport in SNs before and after Brefeldin A (BFA) treatment analyzed by kymograph. The number of neurons analyzed in the experiment is indicated in the bar graphs. Error bars show SEMs. NS, nonsignificant; **p < 0.01, ***p < 0.001. Student’s unpaired two-tailed t-test. **D-E.** Bar graphs show flux and velocity of anterograde and retrograde LV **(D)** and mitochondrial **(E)** transport in SNs before and after 3-hour Docosahexaenoic acid (DHA) treatment analyzed by kymograph. The number of neurons analyzed in the experiment is indicated in the bar graphs. Error bars show SEMs. NS, nonsignificant; **p < 0.01, ***p < 0.001. Student’s unpaired two-tailed t-test. **F-G.** Bar graphs show the flux and velocity of anterograde and retrograde LV **(F)** and mitochondrial **(G)** transport in SNs treated with DHA or DMSO (control) for 24 hours analyzed by kymograph. The number of neurons analyzed in the experiment is indicated in the bar graphs. Error bars show SEMs. NS, nonsignificant. Student’s unpaired two-tailed t-test.

**Figure 8.**
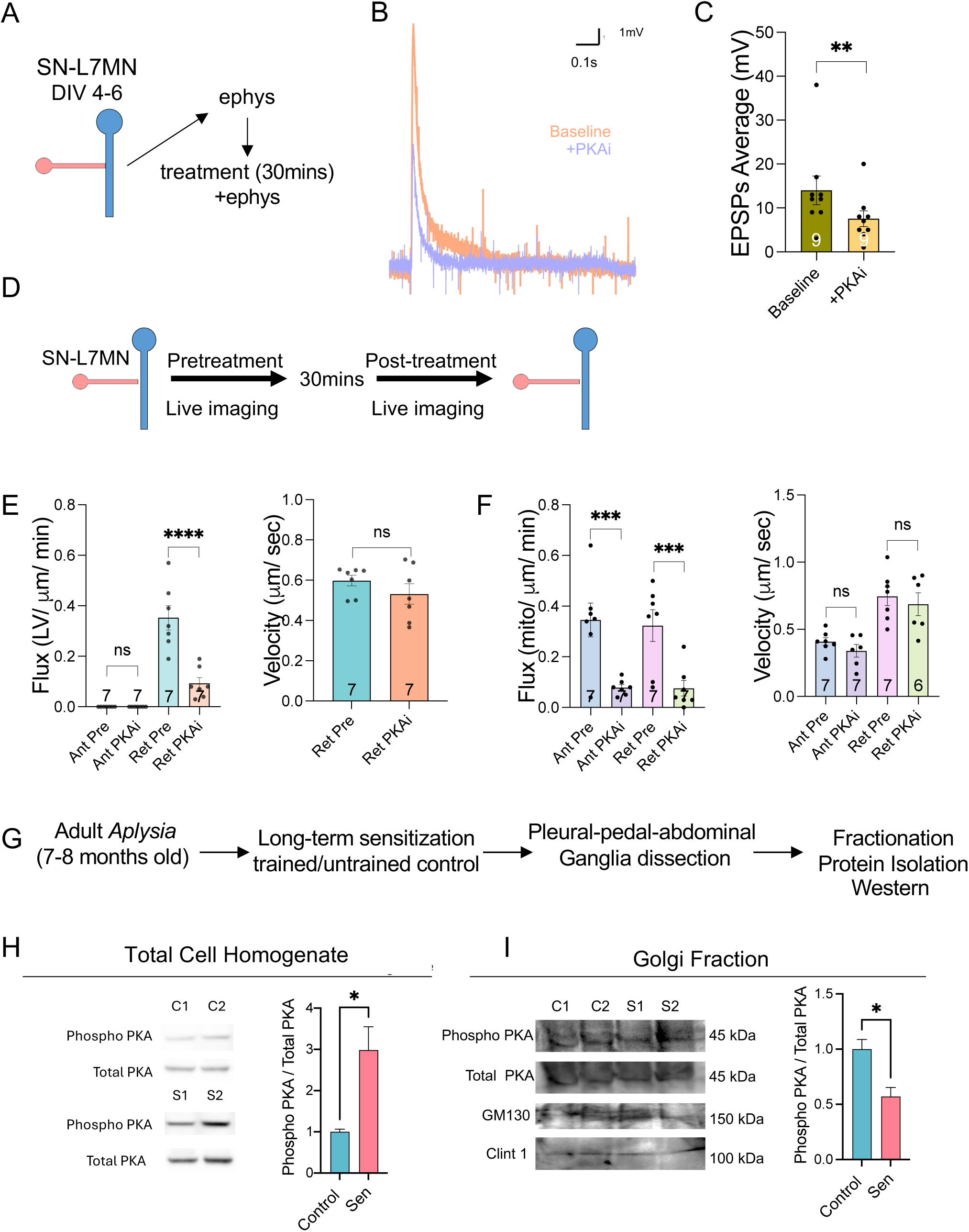
PKA signaling modulates transport of lysosomal vesicles during synapse maintenance. **A.** Experiment design of electrophysiological recordings before and 30 minutes (mins) after pharmacological application. **B.** Representative trace of SNL7MN (DIV 4-6) excitatory postsynaptic potentials (EPSPs) before (pink) and after (purple) 14-22 amide (+PKAi) in millivolts (mV); scale display 1mV along the y-axis and 0.1 seconds (s) in the x-axis. **C.** Bar graph shows average EPSPs in mV of SNL7MN before (pre) and 30 mins after +PKAi application. The number of neurons analyzed in the experiment is indicated in the bar graphs. Error bars show SEMs. ***p < 0.001. Student’s paired t-test. **D.** Experiment design. **E-F.** Bar graphs show the flux and velocity of retrograde lysosomal vesicle **(E)** and bidirectional mitochondrial **(F)** transport before and after PKA inhibition with 14-22 amide (PKAi) analyzed from kymographs, respectively. The number of neurons analyzed in the experiment is indicated in the bar graphs. Student’s paired t-test (lysosomal vesicle flux, mitochondrial flux); Student’s unpaired two-tailed t-test (lysosomal vesicle velocity, mitochondrial velocity). **G**. Experimental schematics for the preparation and analysis of golgi enriched fractions following long-term sensitization training. **H**. Representative western blots for phospho PKA (Thr197) and total PKA and quantitation in total cell homogenates and golgi fractions are shown. N= 24 animals for each condition, Student’s unpaired two-tailed t-test, Error bars show SEMs. NS, nonsignificant; *p < 0.05, **p < 0.01, ***p < 0.001, ****p < 0.0001.

We then considered the possibility that synapse maintenance requires the formation and recycling of synaptic vesicles, leading to a demand for membrane materials. This demand might alter membrane homeostasis following synapse formation, resulting in a reduction in retrograde transport of lysosomal vesicles. To test this idea, we exposed SN-L7MN cultures to Docosahexaenoic acid (DHA) (Figure 7A-7B).

DHA is a major polyunsaturated fatty acid crucial for the brain, playing roles in plasma membrane homeostasis, mitochondrial membranes, and the fusion and fission of vesicles (Tanaka et al., 2012; Kim and Chung, 2007). However, like other membrane materials, DHA is limited. We aimed to determine how lysosomal vesicle and mitochondrial transport might be modulated when there is an abundance of DHA. If lysosomal vesicle transport is reduced due to limited membrane materials, then increasing DHA should enhance lysosomal vesicle and mitochondrial transport. Alternatively, as shown by other studies, DHA exposure might enhance glutamate synaptic transmission and growth, potentially reducing retrograde transport of lysosomal vesicles.

We treated SNL7MN with DHA [50 µM] for 3 and 24 hours. At 3 hours, lysosomal vesicle retrograde flux was reduced, and mitochondrial anterograde flux was enhanced (3 hrs. DHA lysosomal vesicle : Flux: pre-DHA SNL7MN retrograde lysosomal vesicle = 2.19 ± 0.155, n = 11; post-DHA SNL7MN retrograde lysosomal vesicle = 1.83 ± 0.08, n = 11; unpaired t-test, p = 0.033; Figure 7E; Table S7; Video S5. 3 hrs. DHA mitos: Flux: pre-DHA SNL7MN retrograde lysosomal vesicle = 1.04 ± 0.08, n = 11; post-DHA SNL7MN retrograde mitos = 1.09 ± 0.09, n = 11; unpaired t-test, p = 0.63; Figure 7F; Table S7; Video S5).

However, 24-hour DHA treatment did not modulate lysosomal vesicle or mitochondrial transport (24 hrs. DHA lysosomal vesicle : Flux: pre-DHA SNL7MN retrograde lysosomal vesicle = 2.05 ± 0.21, n = 14; post-DHA SNL7MN retrograde lysosomal vesicle = 2.52 ± 0.19, n = 17; unpaired t-test, p = 0.11; Figure 7G; Table S7. 24 hrs. DHA mitos: Flux: pre-DHA SNL7MN retrograde lysosomal vesicle = 2.59 ± 0.23, n = 14; post-DHA SNL7MN retrograde mitos = 2.27 ± 0.28, n = 17; unpaired t-test, p = 0.41; Figure 7H; Table S7). Together, these observations support the possibility that DHA induces neuronal growth, leading to reduced retrograde transport of lysosomal vesicles and enhanced anterograde transport of mitochondria. Consistent with these results, we observed similar changes in the transport of lysosomal vesicles and mitochondria induced by exposure to 5HT, which promotes long-term facilitation (LTF) and the growth of new synaptic connections.

### PKA differentially modulates lysosomal vesicle and mitochondrial transport

Inspired by the similarity of results between DHA and 5HT in modulating lysosomal vesicle transport, and given PKA’s critical role in vesicle budding, we next investigated whether modulation of PKA could be a mechanism underlying the regulation of lysosomal vesicle transport. First, found that thirty minutes after PKA inhibition (PKAi) using 14-22 amide significantly reduced synaptic transmission (EPSPs: Baseline EPSPs average = 14 ± 3.26 SEM, n = 9 neurons; post-PKAi EPSPs average = 7.55 ± 1.78 SEM, n = 9 neurons; paired t-test, p = <0.0001; Figure 9A-9C; Table S1I; Video S6). Next, we found lysosomal vesicle retrograde flux was significantly reduced with PKAi treatment (PKAi: Flux: pre-PKAi SNL7MN retrograde lysosomal vesicle = 0.353 ± 0.04 SEM, n = 7 neurons; post-PKAi SNL7MN retrograde lysosomal vesicle = 0.091 ± 0.02 SEM, n = 7 neurons; paired t-test, p = 0.0008; Figure 8E; Table S8; Video S6).

**Figure 9:**
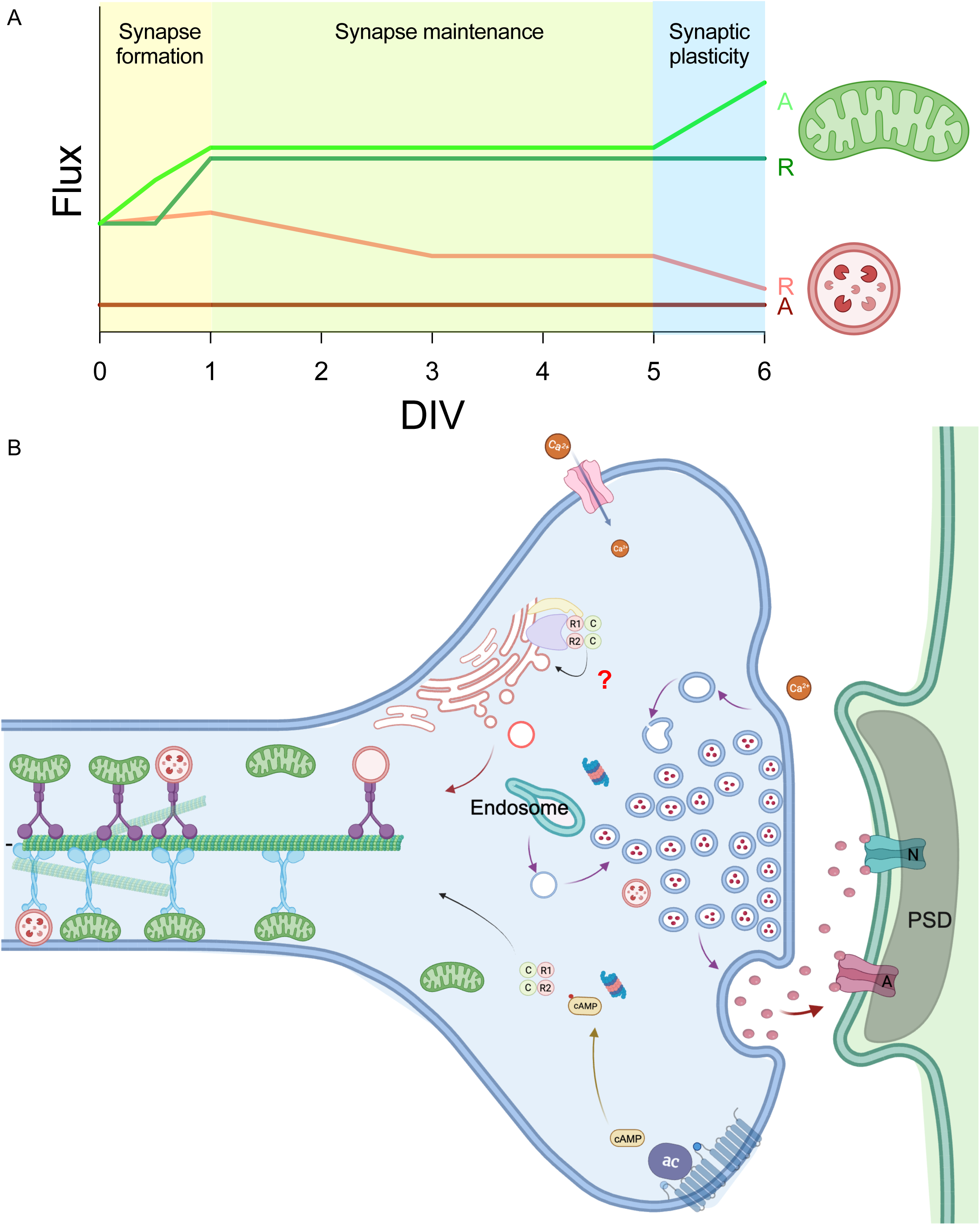
Working model for modulation transport of lysosomal vesicles during synapse maintenance. **A.** Modulation of flux of the bidirectional long-distance transport of mitochondrial and lysosomal vesicles during synapse formation, maintenance, and plasticity. Line drawings indicate changes in transport flux and are based on the measurements described in this manuscript. A: Anterograde, R: Retrograde. Cartoon for organelles are shown. **B.** Proposed model for the reduction in the retrograde transport of lysosomal vesicles during synapse maintenance. Compartmentalization of PKA activity might underlie modulation of mitochondrial and LV transport. Enhancements in PKA activity in the cytoplasm is required for mitochondrial transport. Based on our biochemical experiments, reduction in the activity of PKA in the golgi compartment leads to reduction in LVs. “?” indicates that additional experiments are required to confirm the biochemical findings at a single neuron level.

Additionally, bidirectional transport of mitochondria was significantly decreased with PKAi, corroborating our previous findings that PKA signaling is necessary for modulating mitochondrial transport (PKAi: Flux: pre-PKAi SNL7MN anterograde Mito = 0.345 ± 0.06 SEM, n = 7 neurons; post-PKAi SNL7MN anterograde Mito = 0.077 ± 0.01 SEM, n = 7 neurons; paired t-test, p = 0.008. Pre-PKAi SNL7MN retrograde Mito = 0.324 ± 0.06 SEM, n = 7 neurons; post-PKAi SNL7MN retrograde Mito = 0.074 ± 0.03 SEM, n = 7 neurons; paired t-test, p = 0.003; Figure 8F; Table S8; Video S6). These results suggest that PKA is a critical modulator of the transport of mitochondria and Lysosomal vesicles following synapse formation.

We next considered the possibility that PKA might function in two distinct compartments: in one compartment, PKA activity is enhanced to facilitate mitochondrial transport, while in the other, PKA activity is reduced to limit vesicle budding. Consistent with this hypothesis, compartmentalization of cAMP signaling has been described (Houslay, 2010; Lefkimmiatis and Zaccolo, 2014). To test this, we performed a biochemical fractionation experiment to assess the phosphorylation of PKA. Previous reports indicate that PKA activity on Golgi bodies is critical for the budding of vesicles necessary for lysosome formation (Muñiz et al., 1997) and that protein kinase STK25 inhibits PKA signaling by phosphorylating PRKAR1A (Zhang et al., 2022). We hypothesized that inhibition of PKA activity localized on the Golgi network might reduce the flux of retrograde lysosomal vesicle transport. Importantly, STK25 is also localized to the Golgi network (Preisinger et al., 2004; Matsuki et al., 2010).

To generate sufficient material for biochemical fractionation, we isolated total cell homogenates and Golgi-enriched fractions from the pleural-pedal-cerebral-abdominal ganglia of the central nervous system (CNS) of Aplysia. We first examined whether long-term sensitization (LTS), a form of non-associative learning that produces long-term memory in Aplysia, results in changes in the phosphorylation of PKA at Thr197 (Cauthron et al., 1997). As expected, Western blot analysis of total cell homogenate showed that LTS training resulted in a significant enhancement in Thr197 phosphorylation within one hour of training (Figure 8G&I, N=24 animals, Supplementary Table S8).

We then used a Golgi enrichment kit to prepare Golgi-enriched fractions from the CNS and examined total PKA, phospho-PKA, and markers of the Golgi network, GM130, and Clint1 proteins. Our analysis showed a decrease in phospho-PKA in the Golgi fraction following LTS training (Figure 8G&I, N=24 animals, Supplementary Table S8). Taken together, these results support our notion that compartmentalization of PKA activity is associated with LTS. This suggests the possibility that such modulation of PKA might underlie changes in the bidirectional flux of organelle transport.

## Discussion

The maintenance of synapses once they are formed is a fundamental yet unresolved question in our understanding of brain wiring. While multiple local mechanisms governing synapse maintenance have been identified, the significance of communication between the soma and synapse in maintaining these connections remains poorly understood. Several studies, including our own, have demonstrated that molecular motor-mediated long-distance transport of gene products along microtubules facilitates bidirectional communication between the soma and synapse (Sheetz et al., 1989; Puthanveettil et al., 2008, Swarnkar et al., 2021).

By utilizing Aplysia neuronal cultures, which feature defined synapses between identified neurons that do not form autapses (Kandel, 2001, Alexandrescu and Carew, 2020), we precisely quantified kinestics of the bidirectional long-distance transport in the presence or absence of a functional synapse. Our earlier research revealed that synapse maintenance is associated with persistent enhancements in the bidirectional transport of mitochondria (Badal et al., 2019). Specifically, within 24 hours of synapse formation between sensory neurons (SN) and motor neurons (L7MN), the flux of long-distance anterograde and retrograde transport of mitochondria increased in SN. This finding raised the question of whether the observed enhancements in bidirectional transport are a general phenomenon or specific to mitochondria during synapse maintenance and what mechanisms ensure persistent changes in the long-distance transport for synapse maintenance. To address this, we imaged the bidirectional transport of lysosomal vesicles along with mitochondria during synapse maintenance.

Quantification of lysosomal vesicle transport revealed a significant decrease in the flux of retrograde transport during synapse maintenance, unlike mitochondrial transport (Figure 1). Additionally, the flux of retrograde lysosomal vesicle transport was significantly higher than that of anterograde lysosomal vesicle transport (Figure 1). In contrast, mitochondria exhibited similar anterograde and retrograde transport flux in SN (Badal et al., 2019. These findings illustrate organelle-specific modulation of bidirectional long-distance transport during synapse maintenance. We hypothesized that the increased metabolic demand for maintaining synapses underlies enhanced mitochondrial transport, while the reduction in retrograde transport may be due to more efficient utilization of materials to ensure proper synaptic function, resulting in less waste materials for lysosomal vesicle-mediated processing and recycling. We subsequently tested this hypothesis by examining lysosomal vesicle transport during synapse formation and plasticity.

As previously mentioned, synapse formation is an elegantly orchestrated event involving discrete steps such as target recognition, stabilization, development, and maturation. Anterograde mitochondrial transport significantly increased within 12 hours of plating, became enhanced in both directions within 24 hours, and remained persistent (Badal et al., 2019). This timeframe corresponds to the early stabilization and development of SN-L7MN synapses. During this period, we observed no changes in lysosomal vesicle transport flux; however, at 72 hours, corresponding to the stabilization of excitatory postsynaptic potentials (EPSPs), there was a retrograde-specific reduction in lysosomal vesicle transport flux. We then explored whether synapse formation and maintenance, associated with synaptic plasticity, might produce new changes in bidirectional transport kinetics.

Our observations indicate novel modulation of mitochondrial and lysosomal vesicle transport during synapse formation and maintenance associated with long-term facilitation (LTF) at the SN-L7MN synapses. Specifically, the flux of mitochondrial transport is enhanced in the anterograde direction within an hour of 5×5HT exposure, and this enhancement persists (Figure 3). In contrast, the retrograde transport of lysosomal vesicles is further reduced 24 hours after 5×5HT stimulation (Figure 3). Interestingly, the retrograde transport velocity of lysosomal vesicles and the bidirectional transport velocities of mitochondria show significant enhancements within an hour of 5×5HT stimulation.

The 5×5HT-induced changes in synaptic plasticity and synapse formation serve as a cellular model for learning and LTM. In this model, the formation of new synapses and the remodeling of existing ones are evident by 24 hours after stimulation (Miniaci et al., 2008; Martin et al., 1997). These studies have shown that newly formed synapses can be maintained for several days (Miniaci et al., 2008; Si et al., 2010; Hu et al., 2017). Taken together, these results suggest that lysosomal vesicle transport is modulated for synapse maintenance and that learning produces immediate changes in the kinetics of long-distance transport in an organelle-and direction-specific manner (mitochondria vs. lysosomal vesicles, anterograde vs. retrograde).

Consistent with these observations, previous studies have shown that the molecular motor protein kinesin ApKHC1, which mediates anterograde transport, is upregulated within one hour after 5×5HT stimulation (Puthanveettil et al., 2008). Furthermore, studies have demonstrated that increased metabolic demands associated with plasticity require support from mitochondria anchored close to dendritic spines (Rossi and Pekkurnaz, 2019; Rangaraju et al., 2019; Bapat et al., 2024). Similarly, proper lysosome function is required for synaptic plasticity (Goo et al., 2017; Sun et al., 2022).

Our study focused on the kinetics of long-distance transport, using LysoTracker to stain multiple acidic organelles. Therefore, local lysosomal vesicle function at the synapse could not be elucidated from our measurements. Given that retrograde transport originates from distal neuronal processes, our results suggest an active and persistent mechanism following synapse formation that inhibits the formation of acidic vesicles destined for the neuronal soma. To investigate this further, we conducted experiments to identify signaling pathways underlying the reduction in retrograde transport through pharmacological intervention. A change in transport kinetics suggests a role for modulating transport.

Surprisingly, we found that removing the postsynaptic neuron (L7MN) cell body or inhibiting AMPAR did not alter the transport kinetics of lysosomal vesicles (Figure 4). However, NMDA receptor (NMDAR) inhibition by APV further reduced both anterograde and retrograde transport of lysosomal vesicles, suggesting a role for NMDAR signaling in modulating transport (Figure 4). We cannot exclude the possibility that presynaptic NMDARs, which also play critical roles in synaptic transmission (García-Junco-Clemente et al., 2005; Swarnkar et al., 2018; Lituma et al., 2021), might be involved in modulating transport instead of postsynaptic NMDARs. Since NMDAR activation results in calcium influx, our measurements suggested a role for calcium signaling in modulating transport. Consistent with this assumption, we found that calcium chelation produced a retrograde-specific reduction in the long-distance transport of lysosomal vesicles, unlike the effect of NMDAR inhibition. Intriguingly, long-distance mitochondrial transport was not altered by Ca2+ chelation with BAPTA within 30 minutes of exposure (Figure 4). These results suggest a role for NMDAR and Ca2+ signaling in modulating the retrograde transport of lysosomal vesicles.

Importantly, learning-related signaling pathways such as PKA, PKC, and IP3R did not alter transport kinetics in isolated SNs (Figure S3). Inhibitors of protein degradation altered the transport kinetics of mitochondria without affecting lysosomal vesicle transport, while mTOR inhibition did not alter the transport of either mitochondria or lysosomal vesicles (Figure 5). Autophagy inhibitors bafilomycin and chloroquine specifically impacted retrograde transport kinetics, suggesting a role for autophagy in modulating retrograde transport in isolated SNs (Figure 5). However, in the presence of a synapse, these inhibitors showed organelle-specific modulation (Figure 6). Mitochondrial and lysosomal vesicle transport were significantly decreased, indicating that autophagy regulation during synapse maintenance optimally modulates organelle transport. Synapse maintenance-associated reduction in lysosomal vesicle transport and enhancements in mitochondrial transport depends on optimal levels of autophagy; beyond this, it impairs organelle transport. These observations suggest that local or compartment-specific modulation of signaling pathways might underlie the selective regulation of lysosomal vesicles following synapse formation.

Since lysosomes are generated by vesicles budded off the trans-Golgi network, we considered the possibility that ER-Golgi communication might play a role in modulating lysosomal vesicle transport. However, pharmacological inhibition of ER-Golgi transport by brefeldin A did not alter lysosomal vesicle transport, excluding that possibility. A major change associated with synapse formation in the presynaptic compartment is the development of an active zone and the generation of synaptic vesicles, a process that demands materials for vesicle membranes (Sudhof, 2012). This demand might result in the reduction of retrogradely transported lysosomal vesicles. To test this, we exposed SN-L7MN to DHA, known to increase neuronal growth and synapse function (Figure 7). Intriguingly, we identified modulation of lysosomal vesicle and mitochondria similar to that observed with 5×5HT exposure. A decrease in lysosomal vesicle transport in the retrograde direction and an increase in mitochondrial transport in the anterograde direction within three hours of exposure suggest that these organelles are modulated by changes in neuronal activity.

While these results identified a few key players, we were unable to pinpoint a specific mechanism for the differential modulation of mitochondria and lysosomal vesicles. Our results suggest that modulation of autophagy could indeed be a good candidate mechanism. Several studies have shown that PKA is a key regulator of autophagy (Grisan et al., 2021; Stephan et al., 2009; Overhoff et al., 2022). Our results suggest that inhibition of PKA resulted in the inhibition of mitochondrial transport during synapse maintenance and further reduction in lysosomal vesicle transport (Figure 8). This effect is similar to what we observed with the inhibition of autophagy (Figure 6). Since cAMP/PKA signaling is critical for synapse function, this data can be explained by the compartmentalization of PKA signaling. In one compartment, PKA is inhibited during synapse maintenance, while in another, PKA remains unaffected. The compartmentalization of cAMP signaling has been described (Houslay, 2010; Lefkimmiatis and Zaccolo, 2014). Consistent with this possibility, PKA is known to reside in the trans-Golgi network (Mavillard et al., 2010; Muniz et al., 1997), which is also a source for budding vesicles that form lysosomes. According to a recent study, the regulatory subunit PRKAR1A could be phosphorylated by STK25, resulting in the inhibition of PKA function (Zhang et al., 2022). Interestingly, STK25 also resides on the Golgi (Preisinger et al., 2004; Fidalgo et al., 2010; Matsuki et al., 2010).

Although it is not known whether Golgi-resident STK25 phosphorylates Golgi-resident PKA, resulting in the reduction of vesicle budding from the Golgi for lysosome formation, we considered the possibility that compartmentalized modulation of PKA activity modulates bidirectional long-distance transport of mitochondria and lysosomal vesicles. Modulation of Golgi-resident PKA results in reduced retrograde transport flux, while cytoplasmically localized PKA is critical for mitochondrial transport (Figure 8). Although this model is not proven, our biochemistry data supports compartmentalized regulation of PKA. We found that induction of long-term sensitization, a behavior training that induce robust LTM, resulted in enhancements in PKA phosphorylation at Thr197, as measured using an anti-phospho-Thr197 antibody. Phosphorylation of Thr197 is a physiological mechanism underlying the regulation of PKA (Cauthron et al., 1998). Our attempts to fractionate Golgi networks following long-term sensitization identified a fraction that showed a decrease in Thr197 phosphorylation and a cytosolic fraction that showed an increase in Thr197 phosphorylation. These results support our working model that compartmentalization of PKA activity results in decreased lysosomal vesicle transport and enhanced mitochondrial transport during synapse maintenance (Figure 9). Further work will be required to pinpoint the mechanism underlying compartmentalization at the synapse and regulation following synapse formation.

In summary, our study identifies a previously unrecognized relationship between the long-distance bidirectional transport of lysosomal vesicles and mitochondria and synapse maintenance. Briefly, long-distance transport can be modulated in a direction-specific and organelle-specific manner to fine-tune synapse function. This involves persistent compartmentalization of signaling and modulation of autophagy, resulting in distinct changes in transport kinetics. Given that both mitochondria and lysosomal vesicles require dynein molecular motors for retrograde transport, the modulation should occur at the level of their biogenesis and/or their specific adapter proteins that help them transported by molecular motors. Thus, our findings suggest the possibility of complex regulation of compartmentalized signaling for synapse maintenance. Dissecting the molecular underpinnings of this regulation and understanding how bidirectional communication between the soma and synapse is modified will be necessary for a comprehensive understanding of synapse maintenance.

## Acknowledgements

We gratefully acknowledge funding support from the NSF (Grant 2231247) to SVP, and NIH (1R01MH119541 and 1R01MH118444) to SVP, 1F31MH127958-01A1 to KKB. We sincerely thank Sajid Alam, Malhar Amin, Jenny Dibley, Ania Grodsky, Neha Gopalakrishnan, Bashar Jawich, Lorenzo Thrasher, Nadeen Al-Ostaz, Francesca Oprea, Sydnie Schafer, Olivia Triltsch, and Ashley Ziemer of Michigan State University for their help with data analysis.

## Experimental Methods

### Experimental Model and Subject Details

Sexually mature Aplysia californica (6-9 months old) ranging from sizes of 1-4 grams and 80-120 grams were obtained from the National Resource for Aplysia at the University of Miami’s Rosenstiel School of Marine, Atmospheric, and Earth Science, and were housed in artificial saltwater tanks on-site at the Herbert Wertheim UF Scripps Institute for Biomedical Innovation & Technology. Animals were allowed at least five days to acclimate before being used for neuronal cultures or behavior.

### Aplysia neuronal cultures

Aplysia neuronal cultures were prepared following the protocol described by Montarolo et al. (1986). Briefly, motor neurons were isolated from the abdominal ganglia of Aplysia weighing 1-4 grams, while sensory neurons were isolated from the pleural ganglia of 80-120 grams, 6-month-old Aplysia. The ganglia were digested with dispase II dissolved in modified L15 at 34.5°C (Montarolo et al., 1986; Martin et al., 1997). After digestion, the pleural and abdominal ganglia were rinsed and transferred to a solution consisting of 25% hemolymph (isolated from wild Aplysia) and 75% modified L15 with L-glutamine.

In a silicone-bottom plate, the ganglia were pinned down and de-sheathed. The L7 and L11 motor neurons were then identified, extracted, and plated in a 35mm glass-bottom dish. Sensory neurons were similarly identified, isolated, extracted, and either plated onto motor neurons or isolated on glass-bottom dishes. These cultures were stored at room temperature overnight. The following morning, the neuronal cultures were transferred and incubated at 17-18°C.

### Synapse formation and maturation measurements

In Aplysia in-vitro systems, measurements of electrical connectivity, specifically excitatory postsynaptic potentials, between co-cultured sensory and motor neurons indicate that synapses form as early as 3-12 hours after plating (Coulson & Klein, 1997; Hu et al., 2010). Additionally, presynaptic axonal mitochondrial bidirectional trafficking is enhanced 24 hours after plating (Badal et al., 2019). Consequently, given the dynamic changes within the first 24 hours of plating, days in vitro (DIV) 0-1 are classified as the “synapse formation” stage. Between DIV 2-3, synapses are maturing, referred to as the “synapse maturation” stage (Kim & Martin, 2015). After DIV 3, the majority of the SNL7MN connections are mature and maintained, leading to the classification of DIV 4-6 as the “synapse maintenance” stage (Martin et al., 1997).

### Intracellular electrophysiology measurements

Excitatory postsynaptic potentials (EPSPs) were measured in SNL7MN between DIV 4-6 as described in Miniaci et al. (2008) and Martin et al. (1997). Briefly, a stimulating glass electrode filled with modified L15 media was used to stimulate the presynaptic SN cell body with a 2-millisecond current input. A recording glass electrode, with a resistance of 5-14 Megaohms and filled with 2.3M KCl, was used to impale the postsynaptic L7MN soma and measure membrane potential responses. This setup allowed for the assessment of basal synaptic communication within the SNL7MN circuitry as a result of presynaptic stimulation.

### Microdissection

After electrophysiological confirmation of synapse formation in SNL7MN at DIV 3-4, the L7MN cell body was manually microdissected using a glass electrode to eliminate postsynaptic cell body-generated signals. Following the microdissection, the cultures were rinsed with modified L15 media, and the culture media was replaced with fresh media. The cultures were then incubated at 17.8°C for 24 hours before live imaging.

### Quantitative live imaging of bidirectional transport

One hour before imaging, LysoTracker Deep Red and MitoTracker Green FM were used to fluorescently label live lysosomal vesicles and mitochondria, respectively as we previously described (Badal et al., 2022). Live transport images were collected using a Zeiss 880 confocal microscope at the Max Planck Florida Institute. Axonal regions at least 300 μm from the SN cell body were imaged using a 63x oil objective lens within a 512×512 pixel frame. The imaging parameters were: scan speed of 13 (maximum), scan time of 314 milliseconds, pixel dwell time of 0.51 milliseconds, for 500-600 cycles, with an image size of 45.0 μm x 45.0 μm. A 488 nm laser with a power setting of 1.1 and gain of 560 was used to visualize lysosomal vesicle transport, while a 633 nm laser with a power setting of 1.5 and gain of 650 was used to visualize mitochondrial transport. The live transport images were saved as .czi files.

### Quantitative analysis of time-lapse image data

To analyze transport, movies were separated by channel, rotated, cropped, and kymographs were constructed and analyzed in ImageJ as previously described Miller and Sheetz, 2004, Miller et al., 2005, Baqri et al., 2009. Aplysia axons are complex and thick therefore, a detailed analysis of Aplysia SNs transport is further explained in Badal, et al., 2022. Briefly, we image transport in a segment of the axon between the axon hillock and the axonal terminal. Using ImageJ, we create kymographs of moving particles, draw a line in the center of the kymograph, and count the number of particles that cross the line over time to determine the flux (F) and the slope of lines to determine the speed or velocity (V_INT_) of transported particles. The flux (F) and velocity (V_INT_) are used to determine the concentration or density of moving particles (C_MOV_); F=V_INT_C_MOV_; (Badal et al., 2022). These and other parameters are used to evaluate axonal lysosomal vesicle and mitochondrial transport modulation (Sheetz, 2004, Miller et al., 2005, Baqri et al., 2009; Badal et al., 2022). Averages, standard errors, and significance are calculated based on the data from individual neurons in Excel.

### Pharmacology experiments

One hour before live transport imaging, neurons were treated with LysoTracker Deep Red [75nM] (ThermoFisher Scientific L12492) dissolved in DMSO to visualize lysosomal vesicles, and MitoTracker Green FM [300nM] (ThermoFisher Scientific M7514) dissolved in DMSO to visualize mitochondria. The following pharmacological reagents/vehicles were used to treat SN/SNL7MN: Phorbol 12-Myristate 13-Acetate (PMA) [50nM]/DMSO for 15 mins; Adenophostin A (ADA) [10 μM]/water for 15 mins; Forskolin (FK) [50 μM]/DMSO for 30 mins; 14-22 Amide (PKAi) [360 nM]/water for 30 mins; D(-)-2-Amino-5-phosphonopentanoic acid (AP5) [100 μM] /water for 30 mins; 6-cyano-7-nitroquinoxaline-2,3-dione (CNQX) [50 μM]/water for 15 mins; BAPTA-FM [1mM] for 30mins; Lactacystin (Lact) [50uM] for 30 mins; Chloroquine (CQ) [50uM] for 30 mins; Bafilomycin A (Baf) [10nM] for 30mins; Brefeldin A (BFA) [10ug/mL] for 30 mins; Docosahexaenoic acid (DHA) [50μM] for 3 and 24 hours; actinomycin D (act) [10uM] for 2 hours; serotonin (5HT) [20μM]/ L15 for 5 mins pulses.

### Long-term facilitation and bidirectional transport measurements

Long-term facilitation (LTF) is the cellular plasticity underlying sensitization in Aplysia and has been thoroughly studied to enhance our understanding of learning and memory. Five interspaced pulses of serotonin (5HT) [10-50 μM] induce LTF in the SNL7MN cultures. LTF is characterized by enhanced synaptic transmission, increased synaptic vesicle release, an increased number of varicosity sites, and the upregulation of transcription and translation, all of which contribute to the long-term strengthening of synaptic communication (Bailey & Chen, 1983; Bailey & Chen, 1988; Alberini et al., 1994; Martin et al., 1997; Martin et al., 1997b). To induce LTF in SNL7MN cultures, we applied 20 μM 5HT to the cultures for 5 minutes, followed by a 15-minute rinse, and repeated this 5HT stimulation five times (5×5HT). We measured excitatory postsynaptic potentials (EPSPs) 24 hours later to verify the enhanced EPSPs induced by 5×5HT (Figure S3; Table S3). To investigate mitochondrial and lysosomal vesicle transport during LTF, we imaged transport 1 hour and 24 hours after 5×5HT.

### Long-term sensitization training

To investigate the role of PKA in regulating lysosomal vesicle distribution in the cytosol and on the Golgi-complex membrane, we examined phospho-PKA and PKA protein expression in the Aplysia nervous system after long-term sensitization (LTS) training. Following established sensitization protocols in Aplysia, we trained the animals for LTS as we described recently (Badal et al., 2024), collected the central nervous system 1 hour after training, isolated pleural, pedal and abdominal ganglia from the central nervous system, fractionated the Golgi membrane and cytosolic materials, and conducted western blotting to evaluate PKA protein expression. Long-term sensitization experiments were conducted on 8-month-old (sexually mature), 80-120 gram Aplysia. Each Aplysia was individually housed for five days, with feeding halted 24 hours before the pre-test. On the day of the pre-test, the animals received four siphon touches, and the duration of siphon withdrawal (in seconds) was observed, averaged, and used to group the animals into behavioral groups. On the test day, animals in the sensitization group received four tail shocks, each spaced 30 minutes apart. Each tail shock consisted of four trains, each lasting 1500 milliseconds, with a shock rate of 0.33 pulses per second (PPS). The control group received mock tail shocks. Post-test, four siphon touch withdrawal assessments were performed 1 or 24 hours after LTS or control training before tissue isolation. The pleural, pedal, and abdominal ganglia were collected 1 hour after LTS or control training.

### Golgi analysis and western blotting

We used the Minute Golgi Apparatus Enrichment Kit (Invent Biotechnologies Inc, GO-037) to assess phosphorylated and total PKA in Golgi fractions and homogenates. Briefly, after behavioral training, total homogenates and Golgi fractions were prepared from Aplysia ganglia according to the manufacturer’s protocols. Western blot analysis and protein quantification were performed as previously described (Swarnkar et al., 2021; Espadas et al., 2024).To confirm the presence of Golgi-membrane proteins in the fractions, we used GM-130 (HuaBio, R1608-7) and CLINT1 (Proteintech, 10470-1-AP) as markers. Phospho-PKA alpha/beta (Thr197) Recombinant Polyclonal Antibody (Invitrogen, 711615) and PKA alpha Polyclonal Antibody (Invitrogen, PA5-17626) were employed to detect phosphorylated PKA and total PKA, respectively, in these fractions.

### Quantification and Statistical Analysis

P-values for unpaired two-tailed Student’s t-test, paired two-tailed Student’s t-test, or one-way ANOVA were calculated in either Microsoft Excel or GraphPad Prism. Fisher or Tukey tests were used for post hoc analyses. Significance was defined as p<0.05. The numbers of replications ‘n” and statistical tests used are described in each figure.

## Contacts for Reagents and Resource Sharing

Further information and requests for resources and reagents should be directed to and fulfilled by the Lead Contact, Sathyanarayanan Puthanveettil.

## CONFLICT OF INTEREST STATEMENT

The authors declare no conflicts of interests.

## DATA AVAILABILITY STATEMENT

All experimental data are included in the manuscript.

## AUTHOR CONTRIBUTIONS

SVP and KB designed the project with inputs from KEM; KB carried out all electrophysiology, and the imaging experiments, YZ provided neuronal cultures and helped with illustrations. KEM analyzed all transport data with input from KB, BR and SLV carried out golgi fractionation and western blot experiments. KEM, KB, BLR and SLV interpreted results; KB wrote the paper, and SVP revised the manuscript based on inputs from authors.

## Notes

### Competing Interest Statement

The authors have declared no competing interest.

