## Supplementary Information for "PKA Activity-Driven Modulation of Bidirectional Long-Distance transport of Lysosomal vesicles During Synapse Maintenance"

Abbreviated title: Regulation of bidirectional organelle transport

Key Words: Synapse formation, synapse maintenance, plasticity, long-distance transport, gene expression, lysosome related organelles, mitochondria, Protein kinase A

Number of Supplementary Figures: 3

Number of Supplementary Tables: 8

Number of Supplementary Videos: 6

### Supplementary Figures

Supplementary Figure 1

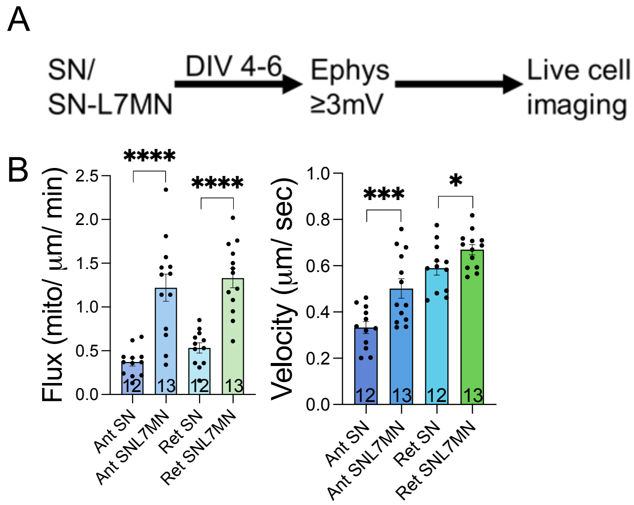


**Supplementary Figure 1. Bidirectional mitochondrial flux and velocity are enhanced during synapse maturation. A.** Experimental strategy for live imaging of organelle transport from SN and SNL7MN cultures after SNL7MN electrophysiology during synapse maintenance. **B.** Bar graphs show average flux and velocity of anterograde (Ant) and retrograde (Ret) mitochondrial transport in SN and SNL7MN 72 hours after synapse formation, *p-Value<0.05, ***p-Value<0.001, ****p-Value<0.0001. Student's unpaired t-test, Error bars are SEM, (see Supplementary Table S2).

Supplementary Figure 2

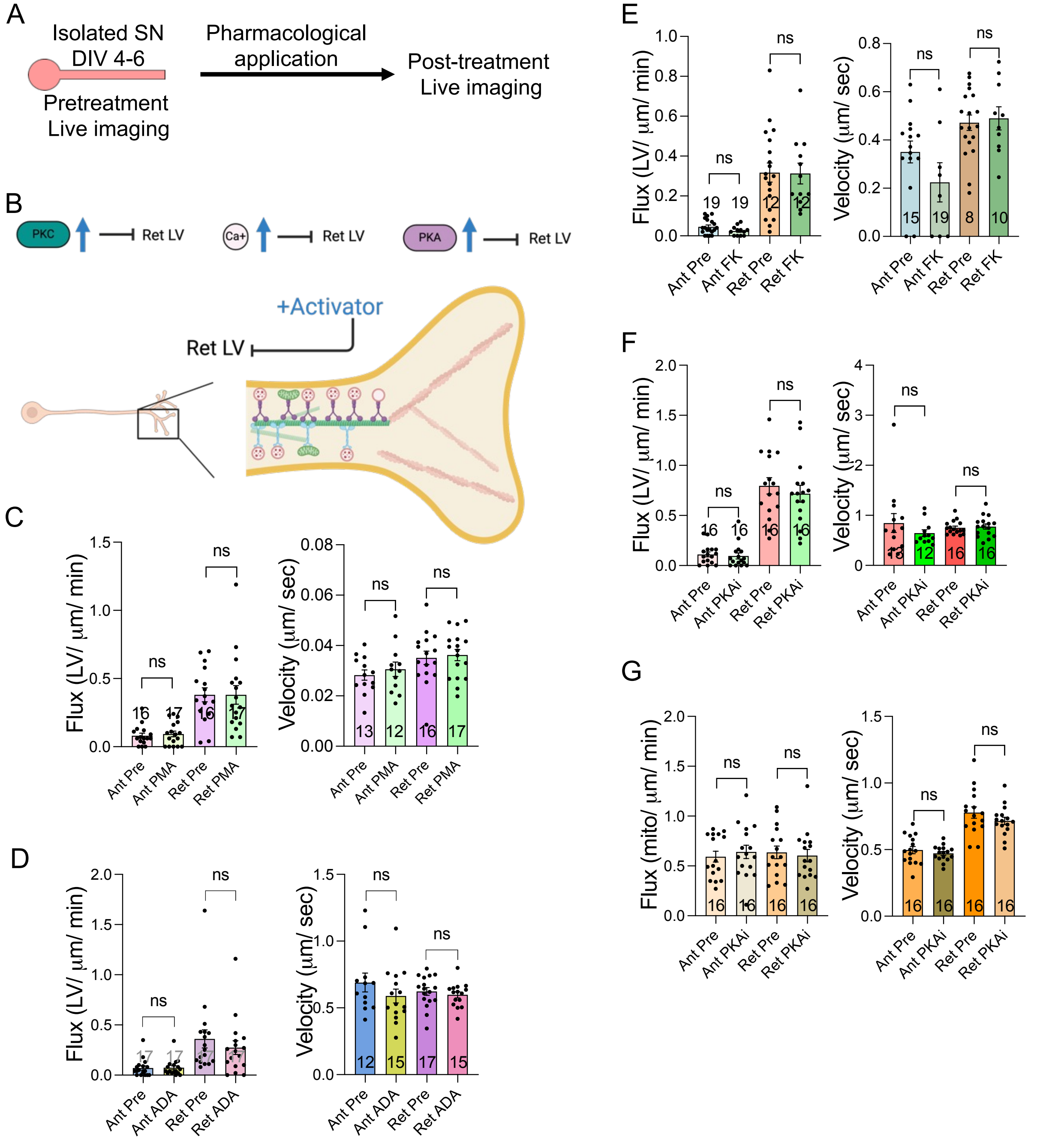


**Supplementary Figure 2. Modulation of reduced LRO transport by learning-related signaling A.** Experiment design. **B.** Cartoon of experimental design and assessing signaling pathways modulating retrograde LV transport in isolated SN. **C-E.** Bar graphs show the flux and velocity of anterograde and retrograde LV transport before and after PKC activation with PMA **(C)**, IP3R-Ca2+ release from ER with ADA **(D)**, or cAMP-PKA activation with Forskolin (FK), **(E)** treatment analyzed from kymographs, respectively. The number of neurons analyzed in the experiment is indicated in the bar graphs. Error bars show SEMs. NS, nonsignificant. Student's unpaired two-tailed t-test (PMA, ADA velocity, and FK velocity); Student's paired t-test (ADA flux, FK flux). **F, G.** Bar graphs show the flux and velocity of anterograde and retrograde LV **(F)**, and mitochondrial **(G)** transport in SNs before and after 14-22 amide (PKAi) treatment analyzed from kymographs. The number of neurons analyzed in the experiment is indicated in the bar graphs. Error bars show SEMs. NS, nonsignificant. Student's paired t-test (flux); Student's unpaired two-tailed t-test (velocity). See supplementary table S5.

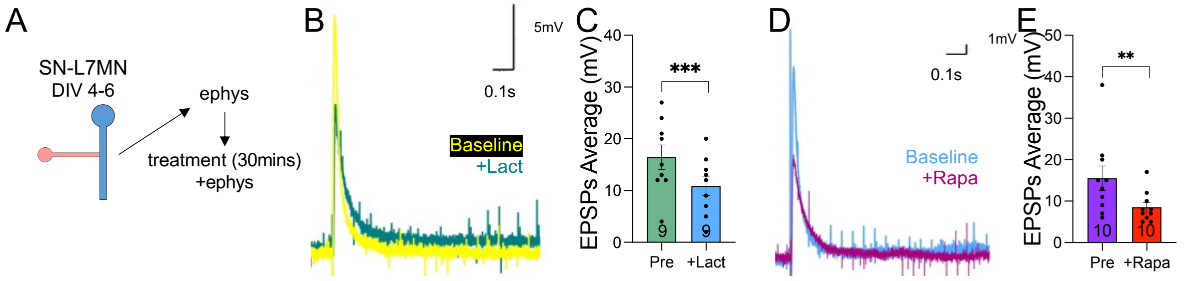


**Supplementary Figure 3. The effect of lactacystin and rapamycin on excitatory post-synaptic potentials. A.** Experiment design of electrophysiological recordings before and 30 minutes (mins) after pharmacological application. **B.** Representative trace of SNL7MN (DIV 4-6) excitatory postsynaptic potentials (EPSPs) before (yellow) and after (green) lactacystin (+Lact) in millivolts (mV); scale display 5mV along the y-axis and 0.1 seconds (s) in the x-axis. **C.** Bar graph shows average EPSPs in mV of SNL7MN before (pre) and 30 mins after +Lact application. The number of neurons analyzed in the experiment is indicated in the bar graphs. Error bars show SEMs. ***$p < 0.001$. Student's paired t-test. **D.** Representative trace of SNL7MN (DIV 4-6) EPSPs before (brown) and after (purple) rapamycin (+Rapa) in mV; scale display 1mV along the y-axis and 0.1 seconds (s) in the x-axis. **E.** Bar graph shows average EPSPs in mV of SNL7MN before (pre) and 30 mins after +Rapa application. The number of neurons analyzed in the experiment is indicated in the bar graphs. Error bars show SEMs. **$p < 0.01$. Student's paired t-test.

### Supplementary table legends

**Supplementary table S1. Overview of reduced retrograde LV flux during the maintenance of SN-L7MN synapses. Supplementary Table related to Figure 1D** shows LV anterograde and retrograde flux and velocity in SN and SNL7MN during synapse maintenance (DIV 4-6). **Supplementary Table related to Figure 1E** shows LV anterograde and retrograde flux and velocity (DIV 4-6) in SN and SNL11MN cultures that do not form functional synapses. **Supplementary Table related to Supplementary Figure S1** shows the mitochondrial anterograde and retrograde flux and velocity in SN and SNL7MN during synapse maintenance (DIV 4-6).

**Supplementary table S2. Retrograde LV flux assessment during synapse formation. Supplementary Table related to Figure 2D** shows LV retrograde flux and velocity in SN and SNL7MN 6 hours after synapse formation. **Supplementary Table related to Figure 2F** shows LV retrograde flux and velocity in SN and SNL7MN 12 hours after synapse formation. **Supplementary Table related to Figure 2H** shows LV retrograde flux and velocity in SN and SNL7MN 24 hours after synapse formation. **Supplementary Table related to Figure 2J** shows LV retrograde flux and velocity in SN and SNL7MN 72 hours after synapse formation.

**Supplementary table S3. LV and mitochondrial transport during long-term facilitation of SN-L7MN synapses. Supplementary Table related to Figure 3C** shows average excitatory postsynaptic potentials (EPSPs) in SNL7MN before and 24 hours after serotonin (5x5HT) long-term facilitation (LTF)-induction. **Supplementary Table related to Figure 3E** shows LV anterograde and retrograde flux in SNL7MN before and 1 hour after 5x5HT-LTF. **Table related to Figure 3F** shows LV anterograde and retrograde velocity in SNL7MN before and 1 hour after 5x5HT-LTF. **Supplementary Table related to Figure 3H** shows mitochondrial anterograde and retrograde flux in SNL7MN before and 1 hour after 5x5HT-LTF. **Table related to Figure 3I** shows mitochondrial anterograde and retrograde velocity in SNL7MN before and 1 hour after 5x5HT-LTF. **Supplementary Table related to Figure 3K** shows LV anterograde and retrograde flux in SNL7MN 24 hour after 5x5L15 (control) or 5x5HT-LTF. **Table related to Figure 3L** shows LV anterograde and retrograde velocity in SNL7MN 24 hour after 5x5L15 (control) or 5x5HT-LTF. **Supplementary Table related to Figure 3N** shows mitochondrial anterograde and retrograde flux in SNL7MN 24 hour after 5x5L15 (control) or 5x5HT-LTF. **Table related to Figure 3O** shows mitochondrial anterograde and retrograde velocity in SNL7MN 24 hour after 5x5L15 (control) or 5x5HT-LTF.

**Supplementary table S4. Assessing the role of the postsynaptic neuron in regulating presynaptic LV transport during synapse maintenance. Supplementary Table related to Figure 4C** shows LV anterograde and retrograde flux and velocity in SNL7MN with in-tact L7MN cell body or in SNL7MN with L7MN cell body removed (CBR). **Supplementary Table related to Figure 4E** shows LV anterograde and retrograde flux and velocity in SNL7MN before and after CNQX application (15mins). **Supplementary Table related to Figure 4F** shows LV anterograde and retrograde flux and velocity in SNL7MN before and after APV application (30mins). **Supplementary Table related to**

**Figure 4H** shows LV anterograde and retrograde flux and velocity in SNL7MN before and after BAPTA application (60mins). **Supplementary Table related to Supplementary Figure 4I** shows mitochondrial anterograde and retrograde flux and velocity in SNL7MN before and after BAPTA application (60mins).

**Supplementary table S5. Assessing the role of protein degradation and autophagy in modulating LV transport in isolated SNs. Supplementary Table related to Figure 5C** shows LV anterograde and retrograde flux and velocity in SN before or 60mins after Lac (lactacystin) application. **Supplementary Table related to Figure 5D** shows mitochondrial anterograde and retrograde flux and velocity in SN before or 60mins after Lac application. **Supplementary Table related to Figure 5E** shows LV anterograde and retrograde flux and velocity in SN before or 60mins after Rapa (Rapamycin) application. **Supplementary Table related to Figure 5F** shows mitochondrial anterograde and retrograde flux and velocity in SN before or 60mins after Rapa application. **Supplementary Table related to Figure 5G** shows LV anterograde and retrograde flux and velocity in SN before or 60mins after CQ (Chloroquine) application. **Supplementary Table related to Figure 5H** shows mitochondrial anterograde and retrograde flux and velocity in SN before or 60mins after CQ application. **Supplementary Table related to Figure 5I** shows LV anterograde and retrograde flux and velocity in SN before or 60mins after Baf (Bafilomycin) application. **Supplementary Table related to Figure 5J** shows mitochondrial anterograde and retrograde flux and velocity in SN before or 60mins after Baf application. **Supplementary Table related to Supplementary Figure 2C** shows LV anterograde and retrograde flux and velocity in SN before or 30mins after PMA application. **Supplementary Table related to Supplementary Figure 2D** shows LV anterograde and retrograde flux and velocity in SN before or 30mins after ADA application. **Supplementary Table related to Supplementary Figure 2E** shows LV anterograde and retrograde flux and velocity in SN before or 30mins after FK (forskolin) application. **Supplementary Table related to Supplementary Figure 2F** shows LV anterograde and retrograde flux and velocity in SN before or 30mins after PKAi (14-22 amide) application. **Supplementary Table related to Supplementary Figure 2G** shows mitochondrial anterograde and retrograde flux and velocity in SN before or 30mins after PKAi (14-22 amide) application.

**Supplementary table S6. Assessing the role of protein degradation and autophagy in modulating LV transport in SNL7MN during synapse maintenance. Supplementary Table related to Figure 6GC** shows average EPSPs in SNL7MN before and 1 hour after CQ application. **Supplementary Table related to Figure 6F** shows LV anterograde and retrograde flux and velocity in SNL7MN before or 60mins after CQ application. **Supplementary Table related to Figure 6G** shows mitochondrial anterograde and retrograde flux and velocity in SNL7MN before or 60mins after CQ application. **Supplementary Table related to Figure 6H** shows LV anterograde and retrograde flux and velocity in SNL7MN before or 60mins after Baf. **Supplementary Table related to Figure 6I** shows mitochondrial anterograde and retrograde flux and velocity in SNL7MN before or 60mins after Baf application. **Supplementary Table related to Figure S4C** shows average EPSPs in SNL7MN before and 1 hour after Lact application.

**Supplementary Table related to Figure S4E** shows average EPSPs in SNL7MN before and 1 hour after Rapa application.

**Supplementary table S7. Organelle-specific modulation of long distance transport by endoplasmic reticulum (ER) to golgi transport and membrane-material availability. Supplementary Table related to Figure 7B** shows LV anterograde and retrograde flux and velocity in SNL7MN before or 60mins after BFA (brefeldin A) application. **Supplementary Table related to Figure 7C** shows mitochondrial anterograde and retrograde flux and velocity in SNL7MN before or 60mins after BFA application. **Supplementary Table related to Figure 7D** shows LV anterograde and retrograde flux and velocity in SNL7MN before or 3 hours after Docosahexaenoic acid (DHA) application. **Supplementary Table related to Figure 7E** shows mitochondrial anterograde and retrograde flux and velocity in SNL7MN before or 3 hours after DHA application. **Supplementary Table related to Figure 7F** shows LV anterograde and retrograde flux and velocity in SNL7MN 24 hours after DMSO (control) or DHA application. **Supplementary Table related to Figure 7G** shows mitochondrial anterograde and retrograde flux and velocity in SNL7MN 24 hours after DMSO or DHA application.

**Supplementary table S8. Assessing the role of PKA modulating LV transport in SNL7MN during synapse maintenance. Supplementary Table related to Figure 8C** shows average EPSPs in SNL7MN before and 1 hour after PKAi application. **Supplementary Table related to Figure 8E** shows LV retrograde flux and velocity in SNL7MN before and 1 hour after PKAi. **Supplementary Table related to Figure 8D** shows mitochondrial anterograde and retrograde flux and velocity in SNL7MN before and 1 hour after PKAi. **Supplementary Table related to Figure 8H and I** show quantification of western blot data. Phosphorylation data normalized to total PKA level.

### Supplementary video legends

**Supplementary videos S1.** Retrograde LV flux in sensory neurons decreases following synapse maintenance in SNL7MN cocultures but is unaffected in SNL11MN cocultures where functional synapses do not form, as shown in Figure 1.

**Supplementary videos S2.** Retrograde LV flux in sensory neurons is not modulated SN compared to SNL7MN within the first 6, 12, and 24 hours after synapse formation but is reduced 72 hours during synapse maturation, related to Figure 2.

**Supplementary videos S3.** Anterograde mitochondrial flux in SNL7MN is increased for LTF, 1 and 24 hours after 5x5HT, related to Figure 3. Retrograde LV flux in SNL7MN is decreased for LTF, 24 hours after 5x5HT, related to Figure 3.

**Supplementary videos S4.** Retrograde LV flux in SNL7MN is decreased for CQ and Baf, related to autophagy and autophagosome-fusion signaling pathways, respectively, related to Figure 6.

**Supplementary videos S5.** Retrograde LV and anterograde mitochondrial flux are modulated for DHA application, related to Figure 7.

**Supplementary videos S6.** Retrograde LV and bidirectional mitochondrial flux are modulated for PKAi application, related to Figure 8.
